# Associations between reproduction-focused life strategy, sex and Borderline Personality Disorder symptom expression: Evidence from the NESARC national study

**DOI:** 10.1101/2025.09.19.676842

**Authors:** Axel Baptista, Bence Csaba Farkas, Nicolas Hoertel, Mark Olfson, Mario Speranza, Pierre O. Jacquet

## Abstract

**Introduction:** Borderline personality disorder (BPD) features span internalizing and externalizing dimensions of psychopathology, and their expression varies across biological sexes. Despite substantial research efforts, major gaps remain in our understanding of this heterogeneity, particularly regarding its developmental underpinnings. Life history theory, a leading framework in evolutionary developmental biology, can help make sense of this heterogeneity.

**Methods:** In a large nationally representative prospective survey (n = 34 653), the National Epidemiologic Survey on Alcohol and Related Conditions (NESARC), we used Multi-group MIMIC models to investigate whether and how the degree to which individuals trade off somatic maintenance against short-term reproductive goals relates to severity of BPD at a general and a single symptom level, and whether these associations differ between men and women in the general adult population.

**Results:** Men and women prioritizing short-term reproductive goals over somatic maintenance were more likely to endorse each of the 9 DSM-IV BPD symptoms through higher BPD severity, and were more likely to express impulsivity (b=0.11; SE, 0.018; *p*<.001) and suicidal/self-mutilation behavior (b =0.24; SE, 0.026; *p*< .001) and less likely to endorse stress-related paranoid ideation (b=-0.066; SE, 0.023; *p*=.004). Finally, females with a reproduction-oriented life history strategy were more likely to endorse affective instability than males (Females: b=0.12; SE, 0.025; *p*<.001; Males: b=-0.015; SE, 0.044; *p*=0.73).

**Conclusions:** An evolutionary-developmental framework, rooted in human life history theory can help make sense of the heterogeneous manifestations of BPD. Our work also opens the door for future research on the interplay of the reproduction/maintenance trade-off with other genetic and environmental factors and developmental processes in explaining the occurrence of BPD symptoms.

## Introduction

Borderline Personality Disorder (BPD) is characterized by a complex symptomatology, including unstable interpersonal relationships, fear of abandonment, identity disturbance, feelings of emptiness, suicidality and self-mutilation, impulsivity, affective instability, inappropriate and intense anger, and stress-related paranoid ideation (1). Clinical studies also indicate that people with BPD are more likely to pursue short-term reproductive goals (2,3) (e.g., compulsive sexuality, early parenting) and neglect somatic maintenance goals (4–6) (e.g., health-damaging behaviors, general medical co-morbidities). These seemingly unrelated manifestations contribute to a reduced psychosocial functioning, well-being and life expectancy.

We recently showed that this cluster of behaviours, which can be understood as a strategy of life whereby people ‘trade’ somatic maintenance against short-term reproduction, is associated with the sum of adverse experiences they made during childhood, and is highly predictive of the presence of a BPD diagnosis (7). This suggests a view of BPD as the consequence of an adaptive response to childhood experiences signalling future risks of death and disease (8). The production of cognitive, emotional, and behavioral traits that facilitate reproduction at an early age could increase the individuals’ chance to pass on their genes despite a prolonged exposure to morbidity and mortality risks.

However, BPD is highly heterogeneous, spanning a wide array of domains, including internalizing and externalizing dimensions of psychopathology (9–11). Symptom expression also varies across biological sexes. Women are more likely than men to endorse ‘suicidal/self-mutilation behavior’, ‘affective instability’ and ‘chronic feelings of emptiness’, and less likely to endorse ‘impulsivity’ and ‘inappropriate and intense anger’ (12–16). Despite substantial research to delineate distinct BPD profiles, little is known about the developmental pathways that explain their emergence.

The present study aims to account for BPD heterogeneity by investigating whether a ‘reproduction-focused’ life strategy is associated with a distinct BPD subtype, and whether this association differs across biological sexes. We argue that, despite observed differences, the symptom profiles presented by women and men both optimize the achievement of a ‘reproduction-focused’ life strategy.

Indeed, evolution has led to major divergences in the reproductive ecology of women and men (17,18), resulting in the selection of sex-specific psychological traits (8,19). For men, high levels of sensation-seeking, disinhibition and impulsivity may facilitate social dominance and intrasexual competition for mates and status (20). For women, high levels of neuroticism, empathy, and rumination may facilitate interpersonal competence, social support and childrearing ability (20,21). High levels of these traits may confer an increased risk for externalizing disorders in men and internalizing disorders in women, which is consistent with the identified sex-specific pattern of BPD symptoms’ expression (12–16).

Three hypotheses were pre-registered on the Open Science Framework (OSF link: https://doi.org/10.17605/OSF.IO/WKZER), and tested on individual data from the National Epidemiological Survey on Alcohol and Related Conditions (NESARC, N=34,653), using Multiple Indicators Multiple Cause (MIMIC) models. These models test for differential item functioning (DIF), which in this context refers to differences in the likelihood of symptom endorsement between sexes and at different levels of reproduction-focused life strategy (14,22). The pre-registered hypotheses were: 1) a ‘reproduction-focused’ strategy will correlate positively with BPD severity, modelled at the symptom level; 2) a ‘reproduction-focused’ strategy will show DIF for BPD symptoms; and 3) the DIF related to a ‘reproduction-focused’ strategy will vary between women and men.

## Methods

This study followed the ‘Strengthening the Reporting of Observational Studies in Epidemiology’ (STROBE) reporting guideline(23).

### Pre-registration and deviations from the initial pre-registration plan

Our hypotheses and methods were preregistered. Since the registration date, two minor modifications were brought to the statistical methods, detailed and justified in the Supplementary Method S2.

#### Sample

The present study was conducted on data from 34,653 civilian noninstitutionalized U.S. residents of the National Epidemiological Survey on Alcohol and Related Conditions (NESARC), aged 18 years and older and who were successfully reinterviewed at Wave 2. Full details of the cohort and sample selection are available in the Supplementary Method S3, Figure S1).

### Ethics

The authors assert that all procedures contributing to this work comply with the ethical standards of the relevant national and institutional committees on human experimentation and with the Helsinki Declaration of 1975, as revised in 2013. The research protocol, including written informed consent procedures, received full human subjects review, and approval from the U.S. Census Bureau and the Office of Management and Budget.

### Borderline Personality Disorder (BPD) severity

In the NESARC wave 2 interview, all participants were asked about lifetime BPD symptoms. These symptoms were assessed using the National Institute on Alcoholism and Alcohol Abuse Alcohol Use Disorder and Associated Disabilities Interview Schedule-IV, DSM-IV version (24). Analyses for the present study focused on the 9 DSM-IV BPD symptoms (Supplementary Method S1): *Frantic efforts to avoid real/imagined abandonment, unstable/intense interpersonal relationships, identity disturbance, Impulsivity, suicidal/self-mutilation behavior, affective instability, chronic feelings of emptiness, inappropriate/intense anger*, and *stress-related paranoid ideation*. In line with previous NESARC studies focusing on BPD, we only included symptoms causing social or occupational dysfunction (14) (Supplementary Method S1, Table S1). A latent factor underlying these 9 symptom criteria and representing BPD severity was modelled (Supplementary Methods S4). Higher latent scores indicate higher general BPD severity.

### Reproduction-maintenance trade-off

In line with our previous work (7), we modelled a latent factor capturing the shared variance of 4 indicators of the respondents’ reproductive goals (number of children, number of marriages, age at 1^st^ sexual intercourse, history of sexually transmitted diseases) and 3 indicators of their somatic maintenance (Body Mass Index (BMI), perceived physical health, metabolic risk factor) (Supplementary Table S1, Supplementary Methods S4). This latent factor is considered indicative of a reproduction-focused life-history trade-off if it is correlated with more children, more marriages, a younger age at 1^st^ sexual intercourse and more sexually transmitted diseases on one hand, and with a higher BMI, a poorer perceived physical health, and more metabolic problems on the other. The reversed pattern is considered indicative of a maintenance-focused trade-off.

### Covariates

All models were adjusted for age and race (White vs non-White [Black, American Indian/Alaska Native, Asian/Native Hawaiian/Other Pacific Islander, or Hispanic])(25). Race was categorized as White vs non-White because members of minority racial and ethnic groups (i.e., non-White) may experience an overall higher level of environmental adversity, which is a potential confounding factor in our models (26–28). All participants were asked to describe their race by selecting one or more categories defined by the investigator.

### Descriptive statistics

We calculated the means or percentages (as appropriate) of all variable, and their standard errors. All summary statistics and tests for between-group comparison of each variable consider the sampling weights and design effects of the NESARC. Descriptive analyses were conducted using the *survey* R package (29), with the R software version 3.6.1, and are summarized in Tables 1 and in Tables S2-S5 of the Supplementary Material.

**Table 1.**
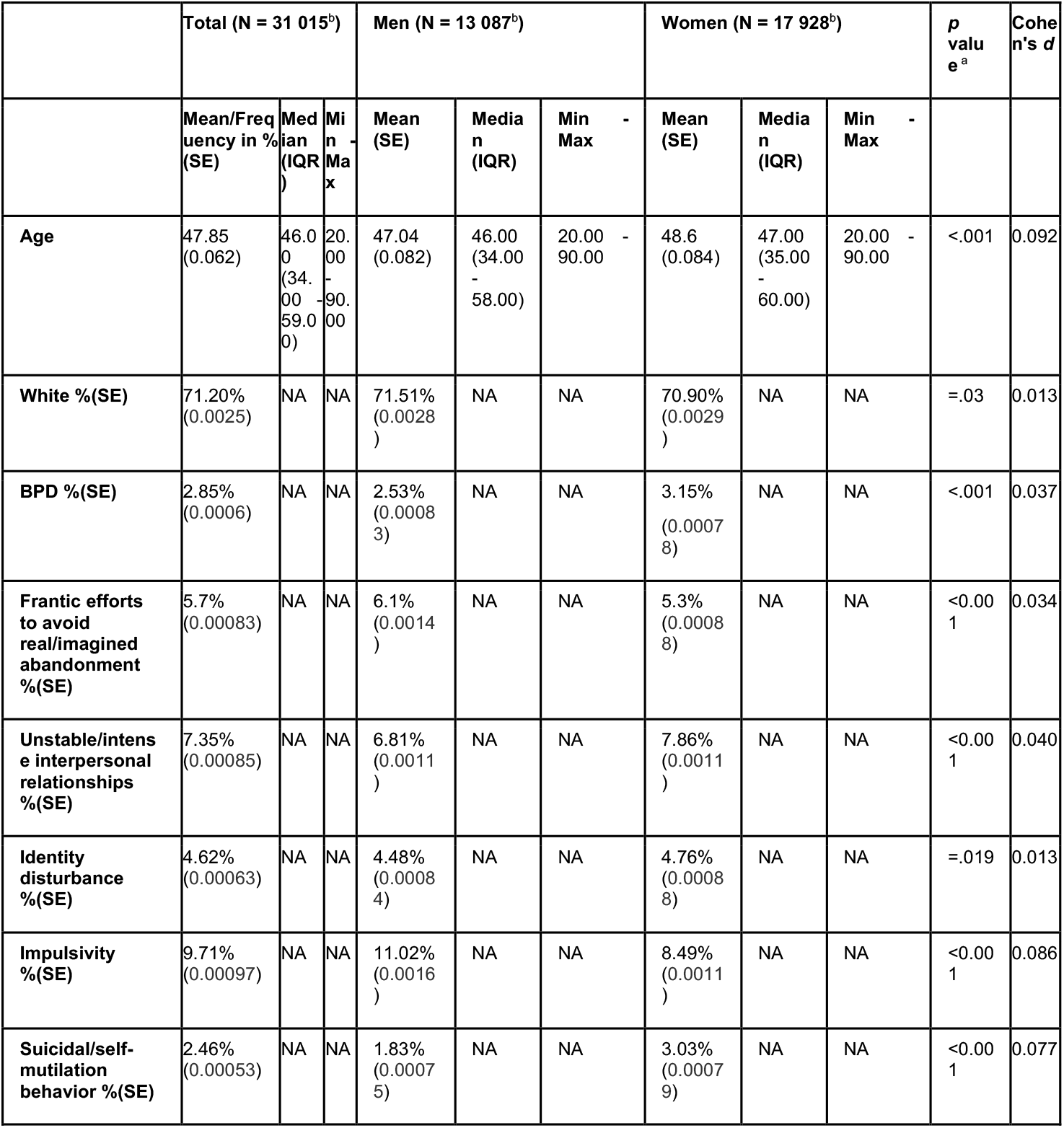
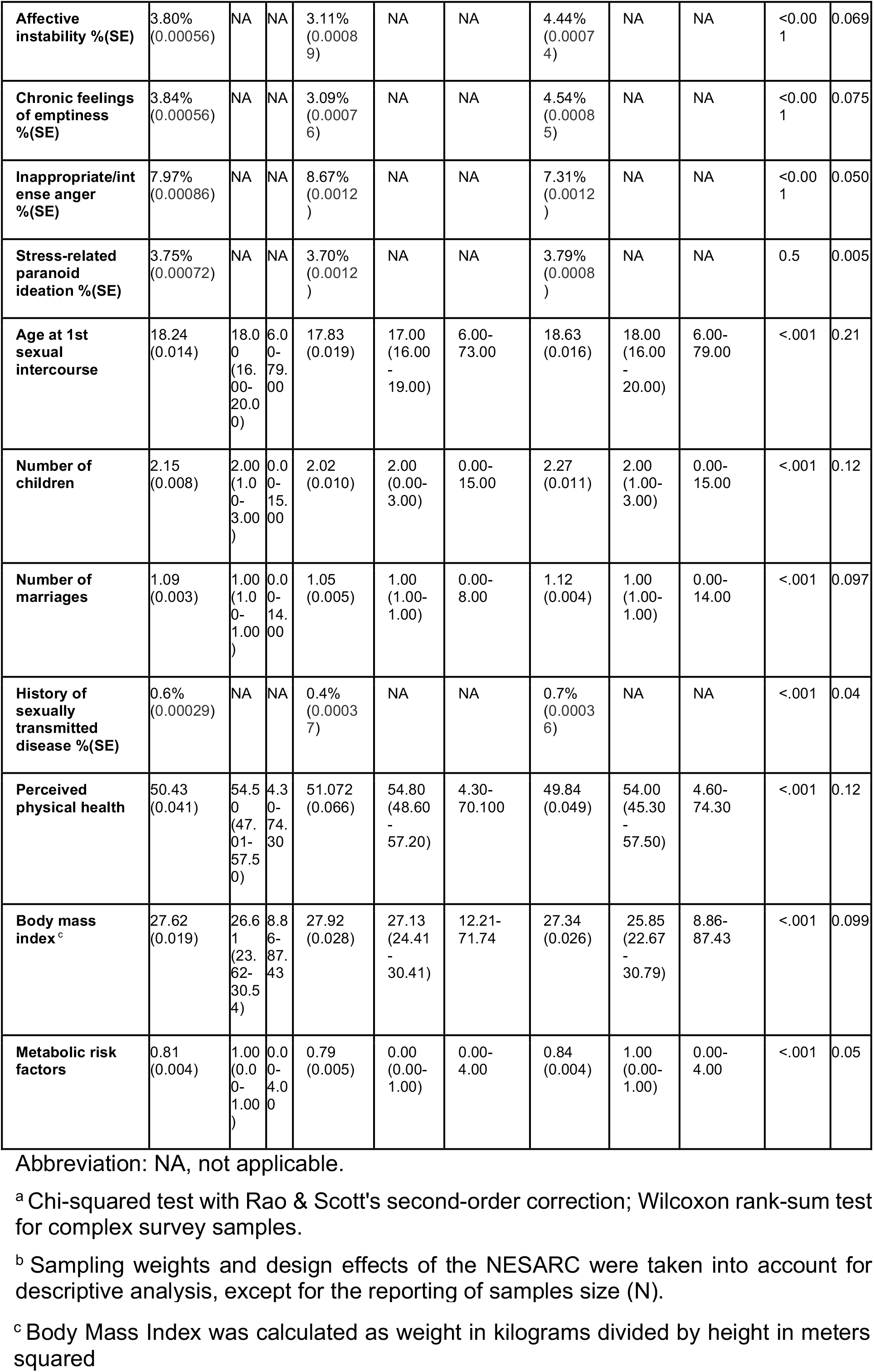
Summary statistics for the variables of interest.

### Multi-group Structural Equation Modelling (SEM)

Multi-group Multiple Indicator Multiple Cause (MIMIC) models from the Structural Equation Modelling (SEM) framework were used as our main multivariate analysis method (22,30) and fitted on the female and male subsamples (Supplementary Methods S4). Models were coded and run in Mplus version 8.1 (31). Parameter estimation was conducted using delta parameterization and the variance-adjusted weighted least squares (WLSMV) estimator, which is appropriate for categorical and dichotomous observed variables and departures from normality (31). Goodness of fit was evaluated using the Root Mean Squared Error of Approximation (RMSEA), Comparative Fit Index (CFI), Tucker-Lewis Index (TLI) and Standardized Root Mean Square Residual (SRMR) statistics. RMSEA values < 0.05, CFI and TLI values > 0.95, and SRMR values < 0.08, which are commonly used to indicate good model fit, were used as cutoffs (32).

For each BPD symptom, DIF between sexes and at different levels of the reproduction-maintenance trade-off was tested through two successive analytic steps (Supplementary Methods S4).

#### Step 1

The structure of the MIMIC model was first identified in a data-driven manner by fitting independent models linking the reproduction-maintenance trade-off latent factor to the BPD severity latent factor and each of the individual symptoms. These models were used to determine: i) whether inclusion of each possible link significantly improved model fit and ii) its equality across sexes, using a Chi-squared difference test for nested models via the DIFFTEST function in the Mplus software (33). An alpha level of .05 was maintained, and *p* values were corrected using the Benjamini-Hochberg procedure.

#### Step 2

The final MIMIC model involved all the links identified at Step 1. This allowed us to examine in a single model the different paths (Supplementary Methods S4) that could generate differences in the probability of observing a BPD symptom including:

- The *direct path* implies a direct effect of the reproduction-maintenance trade-off on the probability of observing a given BPD symptom. A significant direct effect indicates that for 2 individuals with an identical BPD severity but a different reproduction-maintenance trade-off, the probability of observing the symptom is higher in 1 of them. A significant direct effect corresponds to Differential Item Functioning (DIF) that is under the direct dependency of the reproduction-maintenance trade-off. We refer to this direct effect as the ‘reproduction-maintenance trade-off direct DIF’.
- The *indirect path* implies an indirect effect of the reproduction-maintenance trade-off on the probability of observing a given symptom *through* the BPD severity. A significant indirect effect indicates that i) individual differences in the BPD severity contributes to individual differences in the probability of observing a given symptom and that ii) individual differences in the BPD severity is explained by individual differences in the reproduction-maintenance trade-off. In other words, a significant indirect effect corresponds to a DIF that is under the indirect influence of the reproduction-maintenance trade-off *through* the BPD severity. For example, if the probability of observing the symptom ‘impulsivity’ is higher when individuals suffer from a more severe general BPD, which itself is more severe when individuals more strongly favor reproduction over somatic maintenance, then we conclude that reproduction-focused trade-offs contribute, via their effect on the BPD severity, to differences in the symptom ‘impulsivity’. We refer to this indirect effect as the ‘reproduction-maintenance trade-off indirect DIF’.

Based on this model: 1) Hypothesis 1 is confirmed if the association of the reproduction-maintenance trade-off latent factor with the BPD severity latent factor is both statistically significant and positive; 2) Hypothesis 2 is confirmed for a given BPD symptom if there is a reproduction-maintenance trade-off direct or indirect DIF for this symptom; and 3) Hypothesis 3 is confirmed for a given BPD symptom if the reproduction-maintenance trade-off direct or indirect DIF varies between females and males.

Some parameters were allowed to vary between males and females, whereas others were fixed (Supplementary Methods S4) *a priori* based on findings from previous studies reporting sex differences in the modelling of the reproduction-maintenance trade-off (7) and BPD symptoms (14):

### Sensitivity analyses

A sensitivity analysis with an alternative model specification was conducted to examine the robustness of the results obtained at Step 1 and Step 2 (Supplementary Methods S5). This model included a composite score of childhood adversity as a confounding variable, and the above mentioned *a priori* constraints were relaxed. This permitted examination in a single model of sex differences in i) the effect of the general BPD severity on the probability to observe a given symptom; ii) the effect of factors not measured in our main analysis on the probability of observing BPD symptoms.

### k-fold cross-validation

Overfitting risks were estimated by applying a 10-fold cross-validation procedure on the final MIMIC model (34) (Supplementary Methods S6).

## Results

### Sample

The analyses were performed on a sample of 31 015 (89.50% of the initial sample including 13 087 males and 17 928 females), after listwise deletion of participants with missing variables (Table 1).

### Multi-group Structural Equation Model (SEM)

Final MIMIC model showed an excellent fit (CFI=0.985, TLI=0.980, RMSEA= 0.017[0.016–0.018], SRMR=0.038) with robustness confirmed by a 10-fold cross-validation procedure (Supplementary Methods S6, Table S13). The percentages of categorical items and mean values of continuous items in the female and male subsamples are provided in Supplementary Tables S2 & S3. The observed correlation matrix in the female and male subsamples are provided in the Supplementary Tables S4 & S5.

We provide below a synthesis of the final MIMIC model’s results test our three hypotheses (Figures 1-2). A complete reporting of the parameter estimates is available in the Supplementary Table S7.

**Figure 1.**
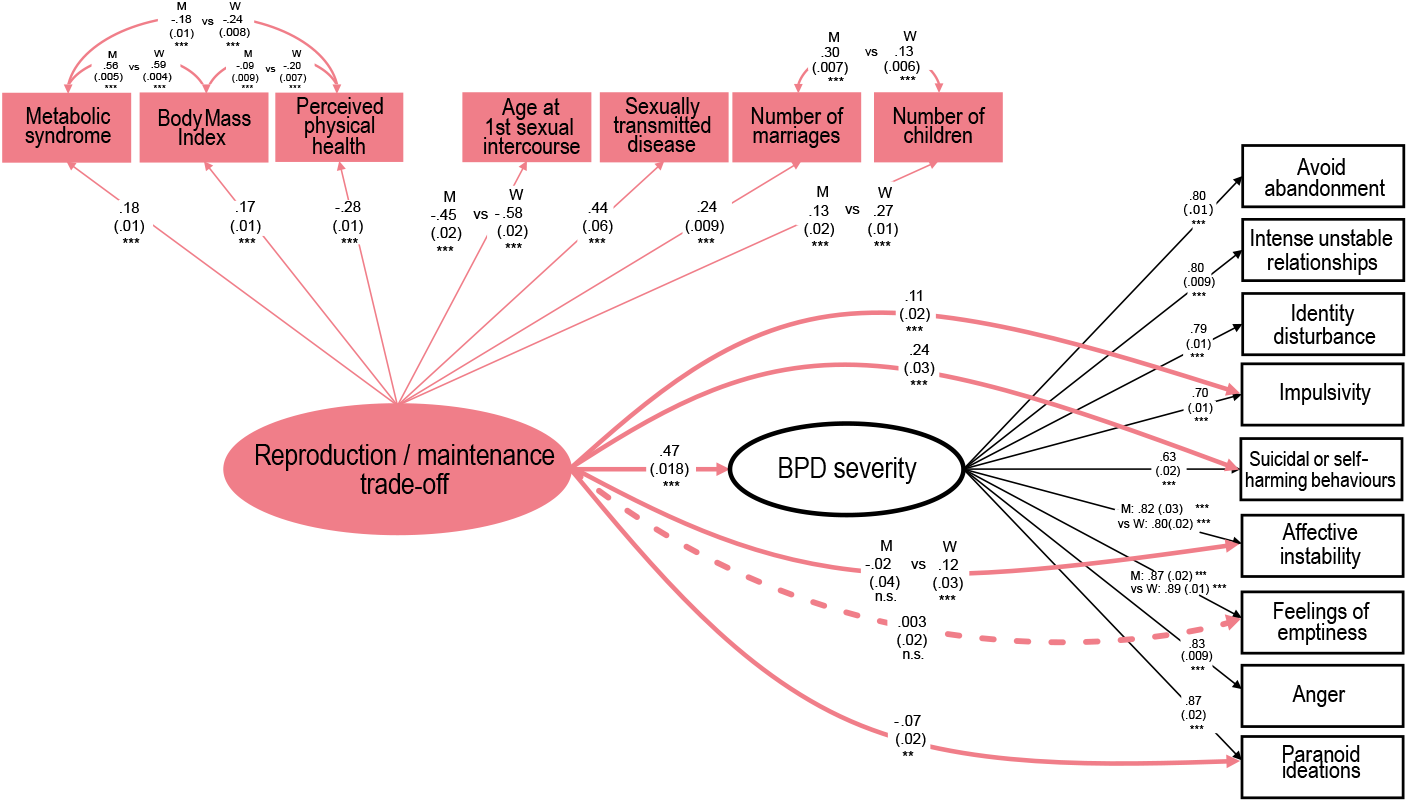
Final Multiple Indicator Multiple Cause (MIMIC) multi-group model in Men and Women. Simplified representation of final MIMIC model. Ellipses represent latent constructs; rectangles represent observed indicators. Paths between indicators and the reproduction/maintenance trade-off latent factor represent factor loadings. Paths between BPD symptoms indicators and the BPD severity latent factor also represent factor loadings. Paths between the reproduction/maintenance trade-off latent factor and BPD symptoms represent direct DIF. Dashed lines represent paths that are not statistically significant. The letter ‘M’ indicates results of the full MIMIC model in the Men subsample. The letter ‘W’ indicates results of the full MIMIC model in the Women subsample. The model was adjusted for age and race. Significance codes: n.s. for p>0.05, ^*^ for p<0.05, ^**^ for p<0.01, ^***^ for p<0.001.

**Figure 2.**
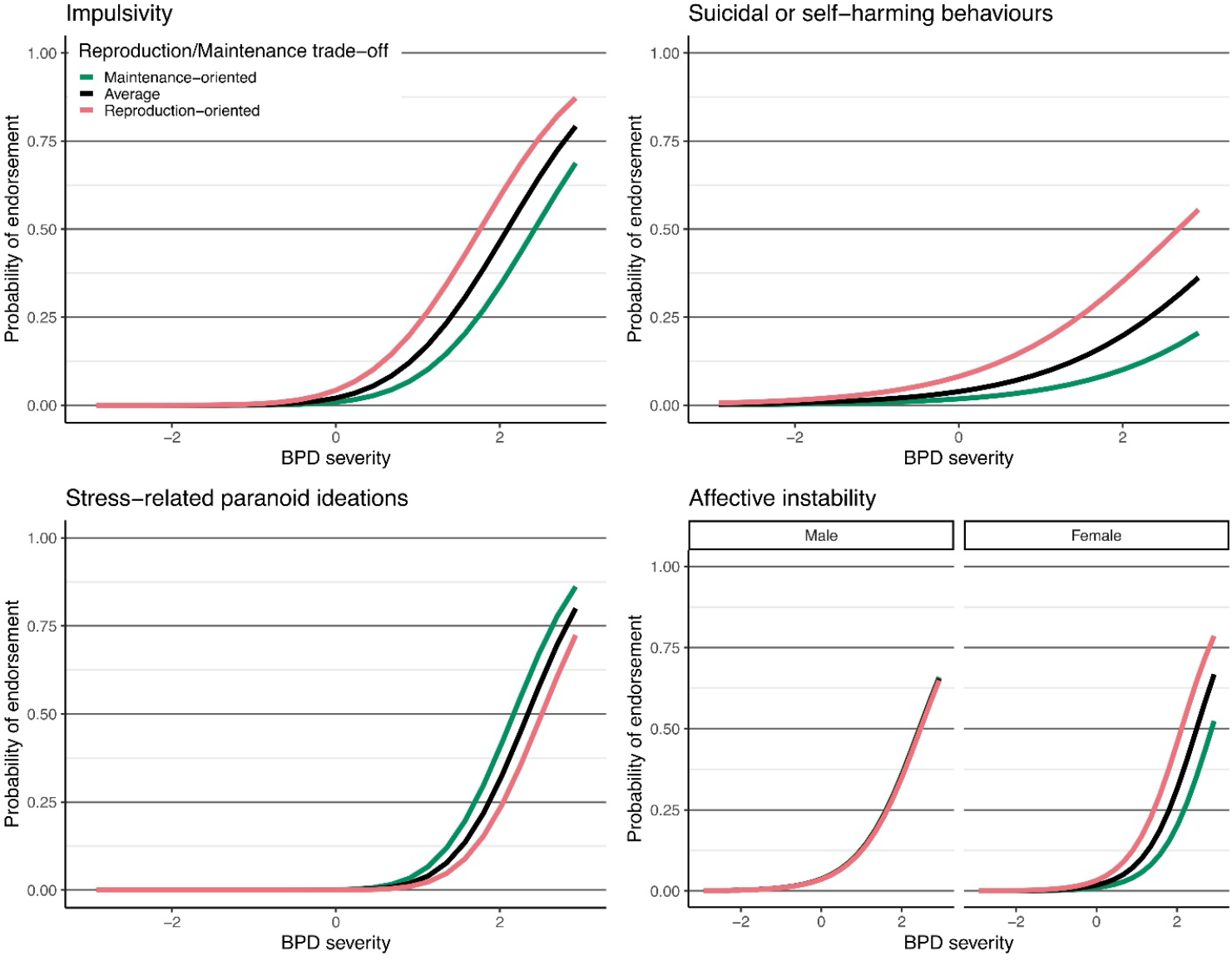
Item Response Curves (IRC), illustrating the 4 identified reproduction/maintenance trade-off related DIF. Illustrations of the differences between average (Black, sample mean), reproduction-focused (Red, +2SD above sample mean) and maintenance-oriented life history strategies (Green, −2SD below sample mean) in the probability of endorsing some BPD symptoms across levels of BPD severity, for the 4 symptoms with significant direct DIF. X axis corresponds to latent BPD severity (in SD from sample mean), y axis corresponds to probability of symptom endorsement. For affective instability, separate subplots show results in the male and female subsamples, as this DIF proved to be sex-dependent.

### Reproduction-maintenance trade-off

Overall, adults who focused more on short-term reproductive goals (more marriages, more children, more sexually transmitted diseases, younger age at 1^st^ sexual intercourse) also presented a poorer somatic maintenance (more metabolic problems, higher BMI, poorer physical health functioning) (Figure 1).

### Hypothesis 1

The model also revealed a positive association between this reproduction-focused trade-off and the latent BPD severity (standardized b=0.46, s.e.=0.016, *p*<0.001).

### Hypothesis 2

A significant direct DIF emerged for 3 symptoms: a positive direct DIF for impulsivity (standardized b=0.11, s.e.=0.018, *p*<0.001) and suicidality/self-mutilation (standardized b=0.24, s.e.=0.026, *p*<0.001); a negative direct DIF for stress-related paranoid ideation (standardized b=-0.066, s.e.=0.023, *p*=0.004). Males and females who focused more on short-term reproductive goals and who exhibited a poorer maintenance were more likely to endorse impulsivity and suicidal/self-mutilation behavior across all levels of BPD severity, and less likely to endorse stress-related paranoid ideation (Figure 2).

There was also a sex-independent indirect effect on the probability of observing the 9 BPD symptoms through BPD severity, as indicated by the significant associations between the reproduction-maintenance trade-off and the BPD severity latent factors (standardized b=0.47, s.e.=0.018, *p*<0.001) as well as between the BPD severity latent factor and each symptom (Figure 1, Table S7). Males and females who focused more on short-term reproductive goals and who exhibited a poorer maintenance had a greater likelihood to endorse each symptom through higher BPD severity.

### Hypothesis 3

The model showed a sex-dependent direct DIF for affective instability (Female: standardized b=0.12, s.e.=0.025, *p*<0.001; Male: standardized b=-0.015, s.e.=0.044, *p*=0.730). Among individuals who focused more on short-term reproductive goals and who exhibited a poorer maintenance, women were more likely than men to express this symptom across levels of BPD severity. A similar sex-dependent pattern was found as a trend for the suicidal/self-mutilation behavior (Table S6).

### Controlling for early life adversity

We ran a sensitivity analysis with an alternative model specification that included childhood adversity as a confounding factor and freely estimated all parameters separately in each sex. It confirmed a positive direct DIF for the impulsivity and the suicidal/self-mutilation behavior symptoms, and a negative direct DIF for the stress-related paranoid ideation. However, it did not confirm a significant direct DIF for the affective instability symptom. It further indicated a significant negative direct DIF for the feelings of emptiness symptom, as well as substantial sex differences in the probability of observing 7 out of the 9 symptoms that were unexplained by factors measured by our main model (Supplementary Methods S5, Tables S8-S12, Figure S2).

## Discussion

According to evolutionary accounts of developmental psychopathology, BPD can be viewed as a consequence of an adaptive response to childhood experiences signalling upcoming risks of death and diseases (8,35). BPD encompasses cognitive, emotional, and behavioral traits that could facilitate reproduction at early age, thereby increasing the individuals’ chance to pass on their genes despite a prolonged exposure to morbidity and mortality risks (36). The present study aimed to account for BPD symptom heterogeneity by investigating whether a ‘reproduction-focused’ life strategy is associated with a distinct BPD profile, and by examining whether this association differs across biological sexes.

The results of our main MIMIC model (Figures 1-2) partly support our hypothesis, showing that i) the extent to which respondents prioritized immediate reproduction over somatic maintenance partly explain individual differences in overall disorder severity; ii) a particular symptom profile was significantly related to reproduction-focused life strategies, beyond the general BPD severity. This profile was characterized by a higher probability of impulsivity, suicidal or self-harming behaviors, and affective instability (in women), with a lower probability of paranoid ideations; and iii) we found evidence for differences in these effects by sex, in that greater probability of affective instability and, as a trend, greater probability of suicidal/self-mutilation behavior, were related to reproduction-focused life history strategies in women but not in men.

This pattern of associations remarkably mirror findings of a recent study of adolescent BPD feature subtypes (37). The three symptoms showing a positive association with reproduction-oriented life histories in our work (impulsivity, suicidal or self-harming behaviors, and affective instability) correspond exactly with the symptoms that characterize the externalized-dysregulation subtype identified in that work, whereas the sole symptom showing a negative association with reproduction-oriented life histories (stress-related dissociation) is a characteristic of the other identified introjective-disturbance subtype. Because that study did not report on sex differences, it remains to future work to investigate whether such BPD subtypes are related to sex in the manner suggested by our results.

Identifying distinct profiles of BPD patients is a necessary step towards personalized medicine that takes into account patient characteristics associated with different treatment responses (38). It is also a necessary step to understanding the etiology of the disorder, as BPD subtypes may result from different developmental trajectories. Despite substantial efforts, there are still major gaps in our understanding of this heterogeneity (38,39). Together with recent theoretical proposals (8,20), we hope that our work will shed light on the functional mechanisms that drive emergence of BPD and sex differences in symptom profiles.

According to sexual selection accounts of psychopathology, sex-invariance is more likely on the externalizing traits of the disorder, such as impulsivity and low harm avoidance. These traits are expected to link more strongly with reproduction-focused life strategies, by their value in securing short-term reproductive benefits in both sexes through intra-sex competition for mates and status (8,40). On the other hand, sex differences are more likely on the internalizing side of the disorder. In this respect, it is remarkable that among people with BPD who prioritized reproduction over somatic maintenance, women exhibited a significantly greater probability of negative emotionality and, as a trend, suicidal/self-mutilation behavior. These traits might contribute to attract attention and empathy from others, a social benefit that could then be exploited by future mothers for childrearing under the form of extra care or resources (20,41). From this perspective, sex differences in BPD symptoms’ expression are not merely seen as purely socially constructed phenomena (14), but rather as evolved psychological traits influenced by environmental and cultural factors (42).

Finally, the lower probability of paranoid ideations among people who favor short-term reproductive goals warrants careful consideration. When severe, these psychotic-like features has been associated with social isolation (43). A plausible explanation for the lower probability of paranoid ideations among individuals pursuing short-term reproductive goals is that they need the opposite: a minimal level of socialization (even if of low quality) to access mating opportunities.

### Limitations

Our study has notable limitations. First, the NESARC sample is representative of the US adult non-institutionalized population. Its findings may not generalize to other populations living in different socio-cultural contexts, or to certain subgroups, such as incarcerated or hospitalized individuals, who have a high prevalence of BPD (1). Second, the small effect size of the direct effects and their differences by sex indicates that the reproduction-focused life strategy construct only explains a small portion of the variance in BPD symptoms. It is possible that some of these symptoms are a consequence of or interact with other symptoms or are related to common mental health comorbidities. Additionally, other genetic and environmental factors, developmental processes, and their complex interactions likely contribute to explaining their occurrence.

### Conclusion

Despite these limitations, our work helps to better understand why BPD symptoms cluster the way they do in some people but not in others. The findings also point to the importance of clinicians viewing BPD symptoms considering patients’ priority motivations and goals, in particular goals related to mating and eliciting care.

## Conflicts of Interest declaration

The authors declare no conflict of interest related to this article.

## Financial Support

A.B. was supported by the Fondation pour la Recherche Médicale (FRM) grant SPF202209015876, and by Fondation de France grant 00112563; P.O.J. was supported by the Agence Nationale de la Recherche grant ANR-22-CE28-0012-01 eLIFUN.

## Data availability statement

In accordance with National Institutes of Health (NIH) guidelines, the data are not publicly available; however, researchers may request specific analyses through the U.S. Census Bureau.

## Notes

https://doi.org/10.17605/OSF.IO/WKZER

### Competing Interest Statement

The authors have declared no competing interest.

