## Supplementary Material for "Associations between reproduction-focused life strategy, sex and Borderline Personality Disorder symptom expression: Evidence from the NESARC national study"

Method S1. LIST OF ITEMS AND SCORING METHODS OF THE VARIABLES INCLUDED IN THE MODELS
Table S1. Items included in the structural equation models (RS = Reverse Coding)

Method S2. DEVIATIONS FROM THE INITIAL PRE-REGISTRATION PLAN

Method S3. SAMPLE DESCRIPTION AND SELECTION
Figure S1. Flowchart of study sample selection

Method S4. MULTI-GROUP STRUCTURAL EQUATION MODELLING (SEM)
Table S2. Univariate higher-order moment descriptive statistics for male
Table S3. Univariate higher-order moment descriptive statistics for female
Table S4. MIMIC model for male: correlation matrix of the variables included
Table S5. MIMIC model for female: correlation matrix of the variables included
Table S6. Data driven identification of the MIMIC model: results of sequential analytic steps
Table S7 - Parameter estimates of the final MIMIC model

Method S5. SENSITIVITY ANALYSIS - INCLUSION OF PARTICIPANTS' EARLY LIFE ADVERSITY AS A CONFOUNDING FACTOR
Table S8. Early life adversity Items (RS = Reverse Coding)

Table S9. Results of the alternative MIMIC model – Results of the sequential analytic steps used to determine the direct effects of the reproduction-maintenance trade-off on BPD symptoms indicators.
Table S10. Results of the alternative MIMIC model – Results of the sequential analytic steps used to determine the variation of the BPD symptoms thresholds between men and women.
Table S11. Results of the alternative MIMIC model – Results of the sequential analytic steps used to determine the variation of the BPD severity indicators slopes between men and women.
Table S12 - Results of the alternative MIMIC model – Parameter estimates of the final alternative MIMIC model
Figure S2. Final alternative Multiple Indicator Multiple Cause (MIMIC) multi-group model in Men vs Women

Method S6. 10-FOLD CROSS VALIDATION
Table S13. 10-fold cross validation

References

### Method S1. LIST OF ITEMS AND SCORING METHODS OF THE VARIABLES INCLUDED IN THE MODELS

**Regarding the measurement of BPD symptoms:** NESARC respondents were asked a series of questions about how they felt or acted most of the time throughout their lives, regardless of the situation or whom they were with (Grant et al., 2008; Harford et al., 2013a, Hoertel 2014). Participants were instructed not to include symptoms that occurred only when they were depressed, manic, anxious, drinking heavily, using medications or drugs, experiencing withdrawal symptoms, or physically ill.

#### Table S1. Items included in the structural equation models (RS = Reverse Coding)

| | | |
|---|---|---|
| Reproduction/maintenance trade-off | reproduction | how old were you when you first had sex/sexual intercourse, or have you never had sexual intercourse |
| | | number of children ever had (incorporating information from wave 1) |
| | | number of marriages (not counting living with someone as if married; incorporating information from wave 1) |
| | | had any other sexually transmitted disease or venereal disease in the past year & did a doctor or health professional confirm the diagnosis |
| | somatic maintenance | body mass index (bmi = reported weight in kilograms/[reported height in centimeters] |
| | | participants' perceived physical health assessed using the physical component summary scale of the widely used 12-item short-form health survey (sf-12) |
| | | z-score of the sum of the following variables: obesity = bmi > 30 (yes/no); had high cholesterol in the previous year (yes/no); had high blood pressure or hypertension in the previous year (yes/no) ; had diabetes or sugar diabetes in the previous year (yes/no) |

| | | |
|---|---|---|
| Borderline personality disorder | 9 DSM-IV BPD symptoms associated with significant distress or impairment | frantic efforts to avoid real/imagined abandonment |
| | | unstable/intense interpersonal relationships |
| | | identity disturbance |
| | | impulsivity |
| | | suicidal/self-mutilation behavior |
| | | affective instability |
| | | chronic feelings of emptiness |
| | | inappropriate/intense anger |
| | | stress-related paranoid ideation |
| covariates | | sex |
| | | age |
| | | ethnicity (white vs. non-white) |

### Method S2. DEVIATIONS FROM THE INITIAL PRE-REGISTRATION PLAN

Deviation 1. According to our pre-registration, our hypotheses are tested at each successive sub-step of the analysis described in Step 1 below (see Method S4). This means that conclusion about the validity of each hypothesis is solely based on the significance of a chi-squared difference test that i) compares a model *with* versus a model *without* the link of interest between the reproduction-maintenance trade-off and a given individual BPD symptom, or that ii) tests the equality of this link between sexes. However, this method does not take into account, for each Step 1 model, the possibility that a part of the variance could be accounted for by some unfitted links between the constructs. Thus, a link identified as significant in one of the Step 1 model may no longer be significant in a more complete model that would include all the links identified as significant at the end of Step 1. This is why we decided to deviate from the initial pre-registration plan and conclude on the validity of our hypotheses based on the final MIMIC model described at Step 2, ie, a model that precisely includes all the links identified as significant at the end of Step 1.

Deviation 2. In our pre-registration we considered the direct paths only, omitting the other paths that generate differences in the probability of observing a BPD symptom. We have decided to deviate from this initial plan by examining, in a single model, the different paths that generate differences in the probability of observing a BPD symptom.

### Method S3. SAMPLE DESCRIPTION AND SELECTION

We used data drawn from the wave 2 of the National Epidemiological Survey on Alcohol and Related Conditions (NESARC). The wave 1 NESARC (2001-2002) is a representative face-to-face survey that includes 43,093 adult residents of households or group quarters in the USA, conducted by the National Institute on Alcoholism and Alcohol Abuse (NIAAA) and described in detail elsewhere (Grant et al., 2009). The wave 2 survey (2004-2005) includes 86.7% of the original sample, corresponding to 34,653 completed interviews. The NESARC wave 2 data were weighted to be representative of the U.S. civilian population based on the 2000 census. The research protocol, including written informed consent procedures, received full human subjects review, and approval from the U.S. Census Bureau and the Office of Management and Budget. The present study was conducted with this sample of 34,653 adults. From this initial sample, participants with missing values in one or several of the observed variables composing the studied dataset were deleted. Analyses were thus performed on 31,015 individual data (89.50% of the initial sample).

### Figure S1. Flowchart of study sample selection

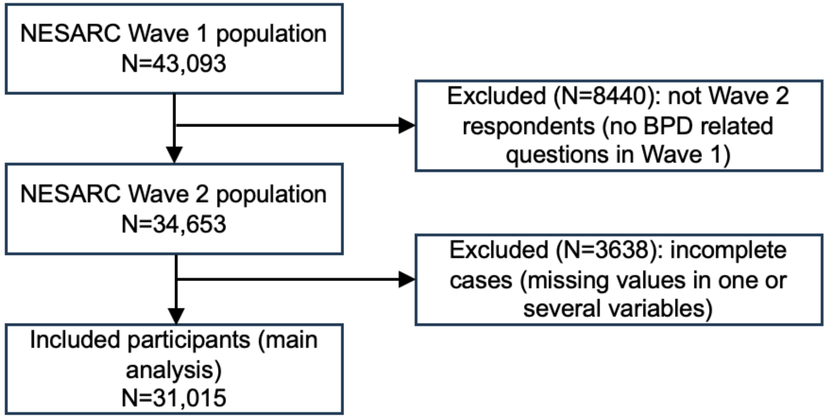

### Method S4. MULTI-GROUP STRUCTURAL EQUATION MODELLING (SEM)

As outlined in the introduction section of the main manuscript, we tested three hypotheses:

H1: A reproduction-focused life history strategy (prioritizing reproductive goals over maintenance ones) would correlate positively with the severity of Borderline Personality Disorder (BPD) (see OSFfig1.jpg for a visual representation of H1), modelled at the symptom level.

H2: A reproduction-focused life history strategy will exhibit differential item functioning (DIF) for BPD symptoms (see OSFfig2.jpg for a visual representation of H2).

H3: The differential item functioning influenced by a reproduction-focused life history strategy would vary between men and women (see OSFfig3.jpg for a visual representation of H3).

To investigate these hypotheses, we implemented Multi-Group Multiple Indicator Multiple Cause (MIMIC) models, a form of Structural Equation Modelling (SEM), on the male and female subsamples (Muthén 1985; Montoya and Jeon 2020). The models were coded and run using Mplus version 8.1 (Muthén and Muthén 2017). We carried out parameter estimation using delta parameterization and the variance-adjusted weighted least squares (WLSMV) estimator, suitable for categorical and dichotomous observed variables and variations from normality (Muthén and Muthén 2017). The goodness of fit was evaluated using the Root Mean Squared Error of Approximation (RMSEA), Comparative Fit Index (CFI), Tucker-Lewis Index (TLI) and Standardized Root Mean Square Residual (SRMR) statistics, with commonly used cut-offs to indicate a good model fit: RMSEA values < 0.05, CFI and TLI values > 0.95, and SRMR values < 0.08 (Hu and Bentler 1999). We compared models using robust Chi squared difference testing via the DIFFTEST feature in the Mplus software, maintaining the usual alpha value of .05. (Asparouhov and Muthén 2010). We corrected p values as necessary using the Benjamini-Hochberg procedure.

#### Each MIMIC model consisted of two key elements

(i) A latent reproduction-maintenance trade-off factor modelled similarly to our preceding study (Baptista et al. 2023), aimed at capturing the shared variance of seven indicators informative of individuals' life history strategies (Nettle et al. 2010; Simpson et al. 2012; Pepper and Nettle 2014; Mell et al. 2018), and listed in the Supplementary Table S1. The participants' reproductive goals were probed by collecting information on the following items: *'number of children ever had'*; *'number of marriages (not counting living with someone as if married)'*; *'how old were you when you first had sex/sexual intercourse, or have you never had sexual intercourse'*; '*history of sexually transmitted disease (confirmed by a doctor or health professional in the year prior the interview)'*. Number of children was self-reported, in response to the following question asked to participants: "How many children have you ever had, including those who are not now living? Please include adopted or foster children and any stepchildren who may have lived with you". Although imperfect, it is the most accurate approximation of the number of biological children available in the database, and thus the best indicator of the respondents' fertility. To compensate for this imperfection, a covariance term was introduced between the indicators 'number of marriages' and 'number of children', the former being more likely to impact the latter with non-biological children than with biological children. Participants' somatic maintenance functions were approximated using the following items: *body mass index* (BMI = reported weight in kilograms/[reported height in centimeters]$^2$); participants' *perceived physical health* assessed using the physical component summary scale of the widely used 12-item Short-Form Health Survey (SF-12, see (Ware Jr et al. 1996)). Their scores were normalized to have a standard deviation of 10 and mean of 50. Lower scores indicate more physical disability. The participants' somatic maintenance state was further assessed by collecting information about their metabolic risk factors. Based on available data in the NESARC database, we approximated participants' *metabolic functioning risk* as the z-scored sum of the following items: (a) obesity (BMI > 30); (b) self-reported past-year presence of high cholesterol; (c) high blood pressure and (d) diabetes, which had to be confirmed by a doctor or health professional. In order to take into account correlation patterns across observed indicators, residual correlations were allowed. For example, we expected that participants who married more often are mechanically more likely to have more children, independently of the resource allocation trade-off latent factor. Similarly, we expected that participants who had higher number of metabolic risk factors and greater BMI would tend to declare a poorer physical health state, independently of that factor. The descriptive statistics of the 7 indicators of the reproduction-maintenance trade-off latent are provided in Tables S2 & S3 right below.

(ii) A latent factor indicating BPD severity modelled using Item Response Theory (IRT) from the 9 indicators that capture the 9 DSM-IV BPD symptoms*: 1-Frantic efforts to avoid real/imagined abandonment; 2-Unstable/intense interpersonal relationships; 3-Identity disturbance; 4-Impulsivity; 5-Suicidal/self-mutilation behavior; 6-Affective instability; 7-Chronic feelings of emptiness; 8-Inappropriate/intense anger; 9-Stress-related paranoid ideation*. The method used corresponds to that employed by Hoertel et al. (2014). The descriptive statistics of the 9 DSM-IV BPD symptoms are provided in Tables 1, S2 & S3. This IRT model describes the relationship between each BPD symptom and BPD severity with a two parameter function (Hoertel et al. 2014). A severity parameter (i.e., the threshold of the link function between BPD severity and BPD symptom) corresponds to the threshold on latent BPD severity factor at which a symptom has a 50% probability of being endorsed. A discrimination parameter (i.e., the slope of the link function between BPD severity and BPD symptom) corresponds to the change in the probability of endorsing the symptom across increasing levels of the latent BPD severity factor. This IRT approach allows for the examination of the probability that a symptom will be observed given a particular level of BPD severity (Hoertel et al. 2014).

**Our analytic plan involved two successive steps that we detail right below:**

**Step 1: this first step comprises three sequential analytic sub-steps used to determine the regression paths that link the two latent constructs, and their variation between men and women**. These sequential analytic sub-steps relied on the iterative comparisons of MIMIC models (Muthén 1985; Montoya and Jeon 2020).

**Sub-step 1.** In an initial sub-step of our analysis, we fitted a multi-group MIMIC model to the male and female subsamples, which included a regression path between the reproduction-maintenance trade-off (see (i) above) and the BPD severity latent factor (see (ii) above).

**Sub-step 2.** In a second analysis sub-step, we examined whether there was a direct effect of the reproduction-maintenance trade-off on each BPD symptoms indicator. For this, we specified a regression path between the reproduction-maintenance trade-off and each BPD symptoms indicator in separates models, supplementing the regression path from the first sub-step of analysis if it proved significant. These paths could vary between sexes. We run a unique model for each of the nine BPD symptom indicators, and compared each of these models with a baseline model that did not include such a regression path. For this comparison, we evaluated the chi-square differences between models using the DIFFTEST procedure (Asparouhov and Muthén 2010). We followed this sequential procedure for all nine items. Given that this method involves 9 Chi-square tests in total, corresponding p values were corrected for multiple comparisons using the Benjamini-Hochberg procedure (Benjamini and Hochberg 1995). All regression paths that were found to be significant through this sequential procedure were then included into a model used for the third and final sub-step of the analysis.

**Sub-step 3.** In a third analysis sub-step, we examined whether the direct effects of the reproduction-maintenance trade-off on BPD symptoms indicators, resulting from the previous analysis sub-step, varied between men and women. For this, we run a MIMIC model which included the significant regression paths derived from the first and second analysis sub-steps. In a baseline model, the direct effects (of the reproduction-maintenance trade-off on BPD symptom indicators, identified from the second sub-step) were constrained to be equal between men and women. We then specified separate alternative models for each BPD symptoms indicator directly associated with the reproduction-maintenance trade-off. In these alternative models, the regression paths between the reproduction-maintenance trade-off and specific BPD symptom were allowed to vary between men and women, while the other direct effects were constrained to be equal between men and women. We compared each of these models with the baseline model which included direct effects that were all constrained to be equal between men and women. For this comparison, we once again evaluated the chi-square differences between the models using the DIFFTEST procedure (Asparouhov and Muthén 2010). The corresponding p values were corrected for multiple comparisons using the Benjamini-Hochberg procedure (Benjamini and Hochberg 1995).

**Step 2: the final MIMIC model involved all the significant parameters identified at step 1 and allows to examine the different (and non-exclusive) paths that generate group differences in the probability of observing a BPD symptom**:

**Path 1:** there is an effect of the BPD severity latent factor on the probability to observe a BPD symptom. This effect is captured by the discrimination parameter (i.e., the slope of the link function between BPD severity latent variable and a BPD symptom, which also corresponds to the loadings of BPD severity indicators). This means that there is variation in BPD severity that contribute to difference in the probability to observe this BPD symptom, above and beyond the effect of the reproduction-maintenance trade-off. For example, if the probability of observing the symptom ‘impulsivity' is all the higher when individuals suffer from a more severe general BPD.

**Path 2 (direct path):** there is a direct effect of the reproduction-maintenance trade-off on the probability of observing a BPD symptom. Such a significant direct effect indicates that, for two individuals with matched latent BPD severity but having different levels of reproduction-maintenance trade-off, the symptom is more likely to be observed in one individual. Consequently, a significant direct effect corresponds to a DIF that directly depends on the reproduction-maintenance trade-off, e.g., if reproduction-focused life strategies are related to impulsivity, irrespective of general BPD severity. This direct effect is now referred to as a reproduction-maintenance trade-off direct DIF. A significant reproduction-maintenance trade-off direct DIF indicates that, for two individuals with an identical BPD severity but a different reproduction-maintenance trade-off, the probability of observing the symptom is higher in one of them.

**Path 3 (indirect path):** there is an indirect effect of the reproduction-maintenance trade-off on the probability to observe a BPD symptom through the BPD severity latent variable. This indirect effect is captured by the joint statistical significance of two estimated associations: (i) between the reproduction-maintenance trade-off and the BPD severity latent factor, (ii) between the BPD severity latent factor and the specific BPD symptom, captured by the loading of the BPD symptom on BPD severity. If both significant, there is evidence for an indirect effect of the reproduction-maintenance trade-off on the probability to observe the specific BPD symptom through the BPD severity latent variable (Fairchild and McDaniel 2017). A significant indirect effect indicates that there is variation in BPD severity that contribute to difference in the probability of observing the BPD symptom, and that this variation is, at least partially, explained by difference in the reproduction-maintenance trade-off. Consequently, a significant indirect effect corresponds to a Differential Item Functioning (DIF) that depends on the reproduction-maintenance trade-off through the BPD severity latent factor. For example, if the probability of observing the symptom ‘impulsivity' is all the higher when individuals suffer from a more severe general BPD, which itself is all the more severe when individuals more strongly favour reproduction over somatic maintenance, then we can conclude that reproduction-focused trade-offs contribute, via their effect on the general BPD severity, to differences in the expression of the symptom ‘impulsivity’. This indirect effect is now referred to as a reproduction-maintenance trade-off indirect DIF.

**Path 4:** the probability to observe a BPD symptom might also be influenced by unknown factors not measured in our model. These influences can be captured altogether in our final MIMIC model by differences in the severity parameters (i.e. the threshold of the link functions between BPD severity and BPD symptoms). For example, if there are sex differences in Impulsivity threshold that are unrelated to the reproduction-maintenance trade-off.

It is important to note that our multi-group approach during step n°1 also allowed us to examine the impact of sex on each of these paths, by comparing a model which includes the path constrained to be equal between men and women with another model which includes the same path free to vary between men and women.

**The final MIMIC model allowed us to validate our hypothesis if:**

**H1** is confirmed if the regression path linking the reproduction-maintenance trade-off with the BPD severity latent factor is both statistically significant and positive.

**H2** is confirmed for a specific BPD symptom if there is a reproduction-maintenance trade-off direct or indirect DIF of this symptom (i.e., path n°2 or n°3).

**H3** is confirmed for a specific BPD symptom if the reproduction-maintenance trade-off direct DIF varies between men and women (i.e., sex-dependent path n°2 or n°3).

Note that we evaluated the strength and direction of the DIF using the standardized regression coefficients (see Table S7) between the reproduction-maintenance trade off and the BPD symptoms indicators, and with the help of Item Response Curves (see Figure 2 in the main manuscript).

**In our main analysis, we integrated the findings from previous studies concerning sex differences in the modelling of the reproduction-maintenance trade-off (by Baptista et al. 2023) and BPD symptoms (by Hoertel et al. 2014)**

Regarding the reproduction-maintenance trade off, the parameters that could vary between men and women included: the loadings of the *'Age at 1st sexual intercourse'* and the *'Number of children'* indicators; the covariance between *the 'BMI'*, the *'Metabolic syndrome'* and the *'Physical health functioning'* indicators. Regarding the BPD severity latent factor modelled using IRT, above and beyond the direct and indirect effects of the reproduction-maintenance trade-off on BPD symptoms: the discrimination parameters (i.e. path n°1) of the *'Affective instability'* and *'Chronic feelings of emptiness'* BPD symptoms as well as the severity parameters (i.e. path n°4) of the *'Suicidal/self-mutilation behavior'*, *'Affective instability'*, *'Chronic feelings of emptiness'* and *'Impulsivity'* BPD symptoms were free to vary between men and women. All other factor loadings and thresholds were fixed to be equal between men and women.

### Table S2. Univariate higher-order moment descriptive statistics for male

| Variable | Sample Size | Mean/Frequency in % | Variance | Skewness | Kurtosis | Minimum | Maximum | % with Min | % with Max | Percentile 20% | Percentile 60% | Percentile 40% | Percentile 80% | Median |
|---|---|---|---|---|---|---|---|---|---|---|---|---|---|---|
| Number of children | 13087 | 2.021 | 3.360 | 1.437 | 4.652 | 0.000 | 15.000 | 26.88% | 0.09% | 0.000 | 2.000 | 1.000 | 3.000 | 2.000 |
| Number of marriages | 13087 | 1.048 | 0.605 | 1.188 | 3.951 | 0.000 | 8.000 | 21.91% | 0.02% | 0.000 | 1.000 | 1.000 | 2.000 | 1.000 |
| Age at 1st sexual intercourse | 13087 | 17.825 | 14.254 | 1.611 | 10.760 | 6.00 | 73.00 | 0.43% | 0.01% | 15.00 | 18.00 | 17.000 | 20.000 | 17.000 |
| Perceived Physical health | 13087 | 51.072 | 97.843 | -1.736 | 2.690 | 4.30 | 70.10 | 0.01% | 0.01% | 45.40 | 55.90 | 52.60 | 57.800 | 54.50 |
| BMI | 13087 | 27.918 | 25.786 | 1.236 | 3.263 | 12.206 | 71.743 | 0.01% | 0.02% | 23.753 | 28.247 | 25.998 | 31.381 | 27.197 |
| Metabolic risk factors | 13087 | 0.788 | 0.943 | 1.180 | 0.786 | 0.000 | 4.000 | 48.90% | 1.60% | 0.000 | 1.000 | 0.000 | 2.000 | 1.000 |
| Age | 13087 | 47.038 | 271.450 | 0.414 | - 0.593 | 20.000 | 90.000 | 0.08% | 0.39% | 33.000 | 51.000 | 42.000 | 64.000 | 47.000 |
| White % | 13087 | 71.51% | | | | | | | | | | | | |
| History of sexually transmitted disease % | 13087 | 0.4% | | | | | | | | | | | | |
| frantic efforts to avoid real/imagined abandonment % | 13087 | 6.1% | | | | | | | | | | | | |
| unstable/intense interpersonal relationships % | 13087 | 6.8% | | | | | | | | | | | | |
| identity disturbance % | 13087 | 4.5% | | | | | | | | | | | | |
| Impulsivity % | 13087 | 11.0% | | | | | | | | | | | | |
| suicidal/self-mutilation behaviour % | 13087 | 1.8% | | | | | | | | | | | | |
| affective instability % | 13087 | 3.1% | | | | | | | | | | | | |
| chronic feelings of emptiness % | 13087 | 3.1% | | | | | | | | | | | | |
| inappropriate/intense anger % | 13087 | 8.7% | | | | | | | | | | | | |
| stress-related paranoid ideation % | 13087 | 3.7% | | | | | | | | | | | | |

### Table S3. Univariate higher-order moment descriptive statistics for female

| Variable | Sample Size | Mean/Frequency in % | Variance | Skewness | Kurtosis | Minimum | Maximum | % with Min | % with Max | Percentile 20% | Percentile 60% | Percentile 40% | Percentile 80% | Median |
|---|---|---|---|---|---|---|---|---|---|---|---|---|---|---|
| Number of children | 18039 | 2.270 | 3.587 | 1.804 | 7.044 | 0.000 | 15.000 | 17.72% | 0.20% | 1.000 | 2.000 | 2.000 | 3.000 | 2.000 |
| Number of marriages | 18039 | 1.123 | 0.589 | 1.790 | 13.322 | 0.000 | 14.000 | 17.95% | 0.01% | 1.000 | 1.000 | 1.000 | 2.000 | 1.000 |
| Age at 1st sexual intercourse | 18039 | 18.627 | 14.786 | 1.879 | 15.163 | 6.00 | 79.00 | 0.50% | 0.01% | 16.00 | 19.00 | 17.000 | 21.000 | 18.000 |
| Perceived Physical health | 18039 | 49.839 | 120.345 | -1.346 | 1.183 | 4.60 | 74.300 | 0.01% | 0.01% | 40.50 | 55.10 | 51.00 | 57.80 | 53.50 |
| BMI | 18039 | 27.337 | 41.228 | 1.322 | 3.523 | 8.857 | 87.428 | 0.01% | 0.01% | 22.299 | 28.290 | 25.018 | 32.594 | 26.571 |
| Metabolic risk factors | 18039 | 0.837 | 1.026 | 1.127 | 0.567 | 0.000 | 4.000 | 46.26% | 2.14% | 0.000 | 1.000 | 0.000 | 2.000 | 1.000 |
| Age | 18039 | 48.589 | 297.985 | 0.392 | -0.702 | 20.000 | 90.000 | 0.05% | 0.81% | 33.000 | 52.000 | 42.000 | 65.000 | 47.000 |
| White % | 18039 | 70.90% | | | | | | | | | | | | |
| History of sexually transmitted disease % | 18039 | 0.7% | | | | | | | | | | | | |
| frantic efforts to avoid real/imagined abandonment % | 18039 | 5.3% | | | | | | | | | | | | |
| unstable/intense interpersonal relationships % | 18039 | 7.9% | | | | | | | | | | | | |
| identity disturbance % | 18039 | 4.8% | | | | | | | | | | | | |
| Impulsivity % | 18039 | 8.5% | | | | | | | | | | | | |
| suicidal/self-mutilation behaviour % | 18039 | 3.0% | | | | | | | | | | | | |
| affective instability % | 18039 | 4.4% | | | | | | | | | | | | |
| chronic feelings of emptiness % | 18039 | 4.5% | | | | | | | | | | | | |
| inappropriate/intense anger % | 18039 | 7.3% | | | | | | | | | | | | |
| stress-related paranoid ideation % | 18039 | 3.8% | | | | | | | | | | | | |

### Table S4. MIMIC model for male: correlation matrix of the variables included

| | 1. | 2. | 3. | 4. | 5. | 6. | 7. | 8. | 9. | 10. | 11. | 12. | 13. | 14. | 15. | 16. |
|---|---|---|---|---|---|---|---|---|---|---|---|---|---|---|---|---|
| 1. Age at 1st sexual intercourse | 1.000 | | | | | | | | | | | | | | | |
| 2. Number of children | -0.098 | 1.000 | | | | | | | | | | | | | | |
| 3. Number of marriages | -0.135 | 0.323 | 1.000 | | | | | | | | | | | | | |
| 4. History of sexually transmitted disease | -0.133 | -0.057 | 0.071 | 1.000 | | | | | | | | | | | | |
| 5. BMI | -0.065 | 0.073 | 0.075 | -0.152 | 1.000 | | | | | | | | | | | |
| 6. Metabolic risk factors | -0.059 | 0.034 | 0.049 | 0.021 | 0.569 | 1.000 | | | | | | | | | | |
| 7. Perceived Physical health | 0.131 | -0.027 | -0.036 | -0.096 | -0.135 | -0.229 | 1.000 | | | | | | | | | |
| 8. Unstable/intense interpersonal relationships | -0.159 | 0.076 | 0.107 | 0.197 | 0.014 | 0.074 | -0.123 | 1.000 | | | | | | | | |
| 9. Identity disturbance | -0.151 | 0.020 | 0.050 | 0.232 | 0.009 | 0.132 | -0.172 | 0.678 | 1.000 | | | | | | | |
| 10. Impulsivity | -0.231 | 0.025 | 0.110 | 0.236 | 0.027 | 0.086 | -0.110 | 0.635 | 0.658 | 1.000 | | | | | | |
| 11. Suicidal/self-mutilation behaviour | -0.226 | 0.080 | 0.075 | 0.220 | 0.010 | 0.118 | -0.182 | 0.521 | 0.492 | 0.565 | 1.000 | | | | | |
| 12. Affective instability | -0.133 | 0.020 | 0.034 | 0.201 | 0.029 | 0.127 | -0.191 | 0.639 | 0.646 | 0.596 | 0.696 | 1.000 | | | | |
| 13. Frantic efforts to avoid real/imagined abandonment | -0.194 | 0.063 | 0.106 | 0.180 | 0.022 | 0.109 | -0.130 | 0.730 | 0.640 | 0.636 | 0.484 | 0.618 | 1.000 | | | |
| 14. Chronic feelings of emptiness | -0.183 | -0.015 | 0.055 | 0.271 | 0.030 | 0.130 | -0.177 | 0.675 | 0.718 | 0.642 | 0.718 | 0.792 | 0.720 | 1.000 | | |
| 15. Inappropriate/intense anger | -0.184 | 0.046 | 0.059 | 0.231 | 0.034 | 0.120 | -0.162 | 0.632 | 0.619 | 0.665 | 0.645 | 0.721 | 0.626 | 0.702 | 1.000 | |
| 16. Stress-related paranoid ideation | -0.159 | 0.020 | 0.048 | 0.200 | -0.001 | 0.097 | -0.172 | 0.639 | 0.696 | 0.621 | 0.580 | 0.705 | 0.659 | 0.735 | 0.759 | 1.000 |

### Table S5. MIMIC model for female: correlation matrix of the variables included

| | 1. | 2. | 3. | 4. | 5. | 6. | 7. | 8. | 9. | 10. | 11. | 12. | 13. | 14. | 15. | 16. |
|---|---|---|---|---|---|---|---|---|---|---|---|---|---|---|---|---|
| 1. Age at 1st sexual intercourse | 1.000 | | | | | | | | | | | | | | | |
| 2. Number of children | -0.212 | 1.000 | | | | | | | | | | | | | | |
| 3. Number of marriages | -0.180 | 0.192 | 1.000 | | | | | | | | | | | | | |
| 4. History of sexually transmitted disease | -0.188 | 0.025 | 0.046 | 1.000 | | | | | | | | | | | | |
| 5. BMI | -0.072 | 0.086 | 0.003 | -0.029 | 1.000 | | | | | | | | | | | |
| 6. Metabolic risk factors | -0.079 | 0.056 | 0.009 | 0.016 | 0.600 | 1.000 | | | | | | | | | | |
| 7. Perceived Physical health | 0.117 | -0.063 | -0.034 | -0.108 | -0.229 | -0.282 | 1.000 | | | | | | | | | |
| 8. Unstable/intense interpersonal relationships | -0.224 | 0.056 | 0.080 | 0.237 | 0.056 | 0.101 | -0.137 | 1.000 | | | | | | | | |
| 9. Identity disturbance | -0.189 | 0.039 | 0.092 | 0.249 | 0.082 | 0.102 | -0.130 | 0.720 | 1.000 | | | | | | | |
| 10. Impulsivity | -0.267 | 0.047 | 0.110 | 0.357 | 0.102 | 0.115 | -0.144 | 0.628 | 0.604 | 1.000 | | | | | | |
| 11. Suicidal/self-mutilation behaviour | -0.311 | 0.073 | 0.188 | 0.313 | 0.097 | 0.117 | -0.220 | 0.548 | 0.506 | 0.592 | 1.000 | | | | | |
| 12. Affective instability | -0.286 | 0.073 | 0.118 | 0.325 | 0.106 | 0.163 | -0.205 | 0.667 | 0.717 | 0.643 | 0.724 | 1.000 | | | | |
| 13. Frantic efforts to avoid real/imagined abandonment | -0.220 | 0.018 | 0.119 | 0.217 | 0.107 | 0.143 | -0.153 | 0.724 | 0.645 | 0.636 | 0.584 | 0.661 | 1.000 | | | |
| 14. Chronic feelings of emptiness | -0.226 | 0.058 | 0.097 | 0.305 | 0.067 | 0.114 | -0.192 | 0.720 | 0.734 | 0.644 | 0.751 | 0.832 | 0.713 | 1.000 | | |
| 15. Inappropriate/intense anger | -0.214 | 0.061 | 0.090 | 0.293 | 0.101 | 0.129 | -0.162 | 0.650 | 0.652 | 0.630 | 0.678 | 0.771 | 0.644 | 0.778 | 1.000 | |
| 16. Stress-related paranoid ideation | -0.172 | 0.048 | 0.104 | 0.300 | 0.097 | 0.133 | -0.140 | 0.661 | 0.664 | 0.605 | 0.623 | 0.741 | 0.665 | 0.753 | 0.752 | 1.000 |

### Table S6. Data driven identification of the MIMIC model: results of sequential analytic steps.

| | Test for the existence of a Reproduction-maintenance trade-off – BPD symptom direct link (analytic sub-step n°2) | | Test for the variation between men and women of the Reproduction-maintenance trade-off – BPD symptom direct link (analytic sub-step n°3) | |
|---|---|---|---|---|
| Symptom | χ2 difference [a] | p difference (Uncorr / Benjamini-Hochberg) | χ2 difference [b] | p difference (Uncorr / Benjamini-Hochberg) |
| Frantic efforts to avoid real/imagined abandonment | 2.628 | 0.2688/ 0.30240 | | |
| Unstable/intense interpersonal relationships | 4.035 | 0.1330/0.171 00 | | |
| Identity disturbance | 5.425 | 0.0664/0.099 60 | | |
| Impulsivity | 17.237 | 0.0002/0.000 6 | 0.743 | 0.3886/ 0.6476667 |
| Suicidal/self-mutilation behavior | 52.624 | < 0.001/< 0.001 | 4.834 | 0.0279/ 0.0697500 |
| Affective instability | 10.502 | 0.0052/0.011 70 | 6.912 | 0.0086/ 0.043000 |

| | | | | |
|---|---|---|---|---|
| Chronic feelings of emptiness | 7.710 | 0.0212/0.038 16 | 0.034 | 0.8538/ 0.8538000 |
| Inappropriate/intense anger | 2.233 | 0.3274/0.327 40 | | |
| Stress-related paranoid ideation | 16.730 | 0.0002/0.000 6 | 0.263 | 0.6082/ 0.7602500 |

Results of sequential analytic sub-steps n°2 and 3 used to determine the regression paths that link the reproduction-maintenance trade-off and BPD symptoms, and their variation between men and women.

[a] chi-square differences between a model with versus a model without the link of interest between the reproduction-maintenance trade-off and the individual BPD symptom of interest, using the DIFFTEST procedure (Asparouhov and Muthén 2010). A significant difference, highlighted in colour, indicates that there is a direct effect of the reproduction-maintenance trade-off on the probability of observing the BPD symptom.

[b] chi-square differences between (i) a model in which the direct effect of interest (of the reproduction-maintenance trade-off on the BPD symptom indicator) is constrained to be equal between men and women, and (ii) a model in which this direct effect is allowed to vary between men and women, while the other direct effects are constrained to be equal between men and women. A significant difference, highlighted in green, indicate that the direct effect of the reproduction-maintenance trade-off on BPD symptoms indicator, resulting from the previous analysis sub-step [a], varies between men and women.

### Table S7 - Parameter estimates of the final MIMIC model.

| **Final Multiple Indicator Multiple Cause (MIMIC) Male model** | | | | | | | |
|---|---|---|---|---|---|---|---|
| **Model part** | **Latent variables** | | **Indicators** | **Unstd. est.** | **se** | **p** | **Std. est.** |
| Measurement model | BPD severity | BY | Unstable/intense interpersonal relationships | 0.718 | 0.010 | < .001 | STDY 0.804 |
| | BPD severity | BY | Identity disturbance | 0.712 | 0.011 | < .001 | STDY 0.791 |
| | BPD severity | BY | Impulsivity | 0.627 | 0.013 | < .001 | STDY 0.703 |

| | | | | | | |
|---|---|---|---|---|---|---|
| BPD severity | BY | Suicidal/self-mutilation behaviour | 0.562 | 0.020 | < .001 | STDY 0.625 |
| BPD severity | BY | Frantic efforts to avoid real/imagined abandonment | 0.715 | 0.010 | < .001 | STDY 0.798 |
| BPD severity | BY | Inappropriate/intense anger | 0.745 | 0.010 | < .001 | STDY 0.834 |
| BPD severity | BY | Stress-related paranoid ideation | 0.771 | 0.013 | < .001 | STDY 0.866 |
| BPD severity | BY | Affective instability | 0.742 | 0.025 | < .001 | STDY 0.823 |
| BPD severity | BY | Chronic feelings of emptiness | 0.774 | 0.016 | < .001 | STDY 0.867 |
| Reproduction-maintenance trade-off | BY | Metabolic syndrome | 0.184 | 0.011 | < .001 | STDXY 0.184 |
| Reproduction-maintenance trade-off | BY | Body Mass Index | 0.085 | 0.007 | < .001 | STDXY 0.167 |
| Reproduction-maintenance trade-off | BY | Perceived physical health | -0.272 | 0.010 | < .001 | STDXY -0.275 |
| Reproduction-maintenance trade-off | BY | Age at first sexuel intercourse | -0.169 | 0.008 | < .001 | STDXY -0.446 |
| Reproduction-maintenance trade-off | BY | History of sexually transmitted diseases | 0.445 | 0.060 | < .001 | STDY 0.437 |
| Reproduction-maintenance trade-off | BY | Number of marriages | 0.186 | 0.007 | < .001 | STDXY 0.239 |
| Reproduction-maintenance trade-off | BY | Number of children | 0.229 | 0.032 | < .001 | STDXY 0.125 |
| Metabolic syndrome | WITH | Perceived physical health | -0.151 | 0.009 | < .001 | STDXY -0.184 |
| Metabolic syndrome | WITH | Body Mass Index | 0.258 | 0.005 | < .001 | STDXY 0.555 |
| Body Mass Index | WITH | Perceived physical health | -0.040 | 0.004 | < .001 | STDXY -0.091 |

| | | | | | | | |
|---|---|---|---|---|---|---|---|
| | Number of marriages | WITH | Number of children | 0.336 | 0.009 | < .001 | STDXY 0.299 |
| Structural model | BPD severity | ON | Reproduction-maintenance trade-off | 0.527 | 0.026 | < .001 | STDXY 0.466 |
| | Impulsivity | ON | Reproduction-maintenance trade-off | 0.108 | 0.018 | < .001 | STDXY 0.107 |
| | Suicidal/self-mutilation behaviour | ON | Reproduction-maintenance trade-off | 0.246 | 0.026 | < .001 | STDXY 0.242 |
| | Affective instability | ON | Reproduction-maintenance trade-off | -0.016 | 0.045 | 0.894 | STDXY -0.015 |
| | Chronic feelings of emptiness | ON | Reproduction-maintenance trade-off | 0.003 | 0.023 | 0.755 | STDXY 0.003 |
| | Stress-related paranoid ideation | ON | Reproduction-maintenance trade-off | -0.067 | 0.023 | 0.004 | STDXY -0.066 |
| Thresholds | | | Unstable/intense interpersonal relationships | 1.472 | 0.014 | < .001 | STDY 1.460 |
| | | | Identity disturbance | 1.703 | 0.015 | < .001 | STDY 1.675 |
| | | | Impulsivity | 1.233 | 0.017 | < .001 | STDY 1.225 |
| | | | Suicidal/self-mutilation behaviour | 2.124 | 0.035 | < .001 | STDY 2.089 |
| | | | Frantic efforts to avoid real/imagined abandonment | 1.613 | 0.015 | < .001 | STDY 1.594 |
| | | | Inappropriate/intense anger | 1.427 | 0.013 | < .001 | STDY 1.414 |
| | | | Stress-related paranoid ideation | 1.806 | 0.018 | < .001 | STDY 1.793 |
| | | | Affective instability | 1.900 | 0.028 | < .001 | STDY 1.865 |
| | | | Chronic feelings of emptiness | 1.886 | 0.030 | < .001 | STDY 1.867 |

**Final Multiple Indicator Multiple Cause (MIMIC) Female model**

| Model part | Latent variables | | Indicators | Unstd. est. | se | p | Std. est. |
|---|---|---|---|---|---|---|---|
| Measurement model | BPD severity | BY | Unstable/intense interpersonal relationships | 0.718 | 0.010 | < .001 | STDY 0.796 |
| | BPD severity | BY | Identity disturbance | 0.712 | 0.011 | < .001 | STDY 0.796 |
| | BPD severity | BY | Impulsivity | 0.627 | 0.013 | < .001 | STDY 0.694 |
| | BPD severity | BY | Suicidal/self-mutilation behaviour | 0.562 | 0.020 | < .001 | STDY 0.612 |
| | BPD severity | BY | Frantic efforts to avoid real/imagined abandonment | 0.715 | 0.010 | < .001 | STDY 0.785 |
| | BPD severity | BY | Inappropriate/intense anger | 0.745 | 0.009 | < .001 | STDY 0.819 |
| | BPD severity | BY | Stress-related paranoid ideation | 0.771 | 0.013 | < .001 | STDY 0.854 |
| | BPD severity | BY | Affective instability | 0.733 | 0.017 | < .001 | STDY 0.799 |
| | BPD severity | BY | Chronic feelings of emptiness | 0.807 | 0.014 | < .001 | STDY 0.894 |
| | Reproduction-maintenance trade-off | BY | Metabolic syndrome | 0.184 | 0.011 | < .001 | STDXY 0.184 |
| | Reproduction-maintenance trade-off | BY | Body Mass Index | 0.085 | 0.007 | < .001 | STDXY 0.132 |
| | Reproduction-maintenance trade-off | BY | Perceived physical health | -0.272 | 0.010 | < .001 | STDXY -0.248 |
| | Reproduction-maintenance trade-off | BY | Age at first sexuel intercourse | -0.223 | 0.006 | < .001 | STDXY -0.579 |
| | Reproduction-maintenance trade-off | BY | History of sexually transmitted diseases | 0.445 | 0.060 | < .001 | STDY 0.414 |

| | | | | | | | |
|---|---|---|---|---|---|---|---|
| | Reproduction-maintenance trade-off | BY | Number of marriages | 0.186 | 0.007 | < .001 | STDXY 0.242 |
| | Reproduction-maintenance trade-off | BY | Number of children | 0.511 | 0.019 | < .001 | STDXY 0.270 |
| | Metabolic syndrome | WITH | Perceived physical health | -0.209 | 0.008 | < .001 | STDXY -0.241 |
| | Metabolic syndrome | WITH | Body Mass Index | 0.336 | 0.005 | < .001 | STDXY 0.590 |
| | Body Mass Index | WITH | Perceived physical health | -0.123 | 0.005 | < .001 | STDXY -0.202 |
| | Number of marriages | WITH | Number of children | 0.152 | 0.007 | < .001 | STDXY 0.128 |
| Structural model | BPD severity | ON | Reproduction-maintenance trade-off | 0.525 | 0.025 | < .001 | STDXY 0.465 |
| | Impulsivity | ON | Reproduction-maintenance trade-off | 0.108 | 0.018 | < .001 | STDXY 0.106 |
| | Suicidal/self-mutilation behaviour | ON | Reproduction-maintenance trade-off | 0.246 | 0.026 | < .001 | STDXY 0.238 |
| | Affective instability | ON | Reproduction-maintenance trade-off | 0.128 | 0.026 | < .001 | STDXY 0.124 |
| | Chronic feelings of emptiness | ON | Reproduction-maintenance trade-off | 0.003 | 0.023 | 0.882 | STDXY 0.003 |
| | Stress-related paranoid ideation | ON | Reproduction-maintenance trade-off | -0.067 | 0.023 | 0.004 | STDXY -0.066 |
| Thresholds | | | Unstable/intense interpersonal relationships | 1.472 | 0.014 | < .001 | STDY 1.445 |
| | | | Identity disturbance | 1.703 | 0.015 | < .001 | STDY 1.685 |
| | | | Impulsivity | 1.398 | 0.016 | < .001 | STDY 1.372 |
| | | | Suicidal/self-mutilation behaviour | 1.943 | 0.027 | < .001 | STDY 1.873 |

| | | | | |
|---|---|---|---|---|
| Frantic efforts to avoid real/imagined abandonment | 1.613 | 0.015 | < .001 | STDY 1.568 |
| Inappropriate/intense anger | 1.427 | 0.013 | < .001 | STDY 1.389 |
| Stress-related paranoid ideation | 1.806 | 0.018 | < .001 | STDY 1.769 |
| Affective instability | 1.759 | 0.023 | < .001 | STDY 1.699 |
| Chronic feelings of emptiness | 1.725 | 0.022 | < .001 | STDY 1.690 |

Elements of the Mplus model syntax (L. K. Muthén & Muthén, 2017): BY can be read as "is measured by"; ON can be read as “is regressed on”; WITH can be read as “is correlated with”.
STDY corresponds to estimates standardization with respect to the latent dependent variable which is suitable for binary variables. The STDY standardized estimate is interpreted as the change in y in y standard deviation units when x changes from zero to one. The STDXY standardized estimate is interpreted as the change in y in y standard deviation units for a standard deviation change in x.
Thresholds correspond to the severity parameter (i.e., the threshold of the link function between BPD severity and BPD symptom) which corresponds to the threshold on latent BPD severity factor at which a symptom has a 50% probability of being endorsed.

### Method S5. SENSITIVITY ANALYSIS - INCLUSION OF PARTICIPANTS' EARLY LIFE ADVERSITY AS A CONFOUNDING FACTOR

In this replication of our analysis, we made two changes to our initial model described in Method S4.

First, we included a variable capturing Early Life Adversity (ELA) as a confounding factor in our MIMIC models (see right below). The aim was to confirm that the paths and their variations between men and women persisted even after taking into account ELA, which we demonstrated in a previous study to impact the reproduction-maintenance trade off as well as the probability of a BPD diagnosis.

Second, in this model we did not incorporate the findings of previous studies on sex differences in the modelling of the reproduction-maintenance trade off and BPD symptoms. The objective was to examine all potential paths that could produce group differences in the probability of observing a BPD symptom, along with their variation between men and women. This included the paths 1 and 4 (Tables S11 and S12, which sex-dependences were predefined in our initial analysis since we incorporated findings from earlier studies).

**Early life adversity.** We conceptualized early life adversity as a sum of environmental risk factors not necessarily correlated with one another but that all contribute to the cumulative probability of being diagnosed BPD. This is at odds with previous research using data from the NESARC, which have generally focused on the single effect of childhood maltreatment on BPD diagnostic. For example, the death of a parent as a child may be independent from being the victim of maltreatment. Even though they are different in nature and occur independently, such factors could cumulatively contribute to the emergence of BPD at later ages (Del Giudice et al. 2011). In line with this cumulative approach, we summed individual z-scores (scaled from 0.0 to 1.0) obtained on 53 items covering 9 environmental risk factors widely known to contribute to early life adversity levels: *Sexual abuse, Physical abuse, Physical neglect, Verbal abuse, Emotional neglect, Household dysfunction, Caregiver psychopathology*, *Traumatic event*, and *Economic scarcity.*

*Sexual abuse, physical abuse, verbal abuse, physical neglect, emotional neglect***.** Participants responded to 19 items informing about their exposure to physical, sexual, and emotional abuse, as well as physical and emotional neglect (Ruan et al. 2008) before the age of 18 (before the age of 10 for the variable "*how often did parent/caregiver leave you alone or unsupervised*"). These questions were adapted from earlier empirically validated scales (Straus 1979; Bernstein et al. 1994). Response options ranged from *'never'* (1) to *'very often'* (5), except for emotional neglect, which ranged from *'never'* to *'always'* and was reverse coded.

*Household dysfunction.* Participants responded to 4 questions asking whether their male caregiver had ever done any of the following to their female caregiver, before they were 18 years old: *'pushed, grabbed, slapped or threw something at her'*; *'kicked, bit, hit with a fist, or hit her with something hard'*; *'repeatedly hit her for at least a few minutes'*; or *'threatened her with a knife/gun or use a knife/gun to hurt her'*. Response options ranged from *'never'* (1) to *'very often'* (5).

*Caregiver psychopathology.* Participants responded to 6 items asking whether before the age of 18, at least one of their caregiver *'had problems with alcohol'* or *'had problems with drugs'*, *'went to jail or prison'*, *'was treated or hospitalized for a mental illness'*, *'attempted suicide'* or *'committed suicide'*. These questions elicited a binary response (*'Yes'* vs. *'No'*).

*Traumatic event.* Participants responded to 23 items about their experience of traumatic events encompassing exposure to war, natural disasters, sexual violence, physical violence, or terrorist attack. Participants who responded *'yes'* to any of these items, and reported having experienced the corresponding event before age 18, were considered to have been exposed to a traumatic event during their childhood.

*Economic scarcity.* Having experienced economic scarcity during childhood was assessed using the following item: 'Before you were 18 years old, was there ever a time when your family received money from government assistance programs like welfare, food stamps, general assistance, Aid to Families with Dependent Children, or Temporary Assistance for Needy Families?'. This question elicited a binary response ('*Yes*' vs. '*No*').

### Table S8. Early life adversity Items (RS = Reverse Coding)

| | | |
|---|---|---|
| early life adversity | physical abuse | before age 18, how often did parent/caregiver push, grab, shove, slap or hit you |
| | | before age 18, how often did parent/caregiver hit you so hard that you had marks or bruises or were injured |
| | emotional abuse | before age 18, how often did parent/caregiver swear, insult or say hurtful things to you |
| | | before age 18, how often did parent/caregiver threaten to hit you or throw something at you |
| | | before age 18, how often did parent/caregiver make you fear that you would be physically hurt or injured |
| | sexual abuse | before age 18, how often did adult/other person fondle/touch you in sexual way when you didn't want this/were to young to know what was happening |
| | | before age 18, how often did adult/other person have you touch them in sexual way when you didn't want this/were to young to know what was happening |
| | | before age 18, how often did adult/other person attempt sexual intercourse with you when you didn't want this/were to young to know what was happening |
| | | before age 18, how often did adult/other person have sexual intercourse with you when you didn't want this/were to young to know what was happening |

| | | |
|---|---|---|
| | physical neglect | before age 18, how often did parent/caregiver make you do chores that were too difficult or dangerous for someone your age |
| | | how often did parent/caregiver leave you alone or unsupervised before 10 years old |
| | | before age 18, how often did you go without things you needed because a parent/caregiver spent the money on themselves |
| | | before age 18, how often did parent/caregiver make you go hungry or not prepare regular meals |
| | | before age 18, how often did parent/caregiver ignore/fail to get you treatment when you were sick |
| | emotional neglect | before age 18, felt there was someone in family that wanted me to be a success (RS) |
| | | before age 18, felt there was someone in family who helped me feel that I was important or special (RS) |
| | | before age 18, felt that my family was a source of strength and support (RS) |
| | | before age 18, felt that i was part of a close-knit family (RS) |
| | | before age 18, felt that someone in my family believed in me (RS) |
| | caregiver psychopathology | before age 18, parent/other adult living in home was problem drinker/alcoholic |

| | | |
|---|---|---|
| | | before age 18, parent/other adult living in home had similar problems with drugs |
| | | before age 18, parent/other adult living in home went to jail/prison |
| | | before age 18, parent/other adult living in home treated/hospitalized for mental illness |
| | | before age 18, parent/other adult living in home attempted suicide |
| | | before age 18, parent/other adult living in home committed suicide |
| | household dysfunction | before age 18, how often did your father/other adult male push, grab, slap or throw something at your mother |
| | | before age 18, how often did your father/other adult male hit your mother with a fist or with something hard |
| | | before age 18, how often did your father/other adult male repeatedly hit your mother for at least a few minutes |
| | | before age 18, how often did your father/other adult male threaten your mother with a knife/gun or use a knife/gun to hurt her |
| | Childhood financial help | before you were 18 years old, was there ever a time when your family received money from government assistance programs like welfare, food stamps, general assistance, aid to families with dependent children, or temporary assistance for needy families? |

| | | |
|---|---|---|
| | traumatic events (before age 18) | ever in active military combat |
| | | ever serve as peacekeeper/relief worker in war zone/other terrorized area |
| | | ever an unarmed civilian in war/revolution/military coup |
| | | ever a refugee |
| | | ever in serious/life-threatening accident |
| | | ever in serious fire, tornado, flood, earthquake or hurricane |
| | | ever sexually assaulted, molested, raped or experienced unwanted sex |
| | | ever physically attacked/beaten/injured by spouse or romantic partner |
| | | ever physically attacked/beaten/injured by anyone else |
| | | ever kidnapped or held hostage or as a pow |
| | | ever stalked by anyone |
| | | ever mugged, held up or threatened with a weapon |

| | | |
|---|---|---|
| | | ever had someone close to you die in terrorist attack |
| | | ever had someone close to you injured in terrorist attack? |
| | | ever yourself injured in terrorist attack |
| | | ever had someone close to you directly experience a terrorist attack (95.34% missing values - excluded from the analysis) |
| | | ever yourself directly experience a terrorist attack |
| | | ever yourself indirectly experience a terrorist attack, like watching on tv |
| | | other than terrorist attack, ever see someone badly injured/killed or ever unexpectedly see a dead body |
| | | other than terrorist attack, ever have someone close to you die unexpectedly |
| | | ever have someone close to you experience any other serious/life-threatening illness, accident or injury |
| | | someone close to you ever have any other very stressful/traumatic experience |
| | | ever yourself have any other very stressful/traumatic experience |

**Table S9. Results of the alternative MIMIC model – Results of the sequential analytic steps used to determine the direct effects of the reproduction-maintenance trade-off on BPD symptoms indicators.**

| | Test for the existence of a Reproduction-maintenance trade-off – BPD symptom direct link (analytic sub-step n°2) | | Test for the variation between men and women of the Reproduction-maintenance trade-off – BPD symptom direct link (analytic sub-step n°3) | |
|---|---|---|---|---|
| Symptom | χ2 difference [a] | p difference (Uncorr / Benjamini-Hochberg) | χ2 difference [b] | p difference (Uncorr / Benjamini-Hochberg) |
| Frantic efforts to avoid real/imagined abandonment | 2.350 | 0.3089/ 0.3971571 | | |
| Unstable/intense interpersonal relationships | 1.782 | 0.4103/ 0.4615875 | | |
| Identity disturbance | 5.041 | 0.0804/ 0.1206000 | | |
| Impulsivity | 35.463 | < 0.001 / < 0.001 | 0.057 | 0.8107/ 0.9151 |
| Suicidal/self-mutilation behavior | 34.504 | < 0.001 / < 0.001 | 1.979 | 0.1595/ 0.6380 |
| Affective instability | 5.066 | 0.0794/ 0.1206000 | | |

| | | | | |
|---|---|---|---|---|
| Chronic feelings of emptiness | 12.098 | 0.0024/ 0.0054000 | 0.011 | 0.9151/ 0.9151 |
| Inappropriate/intense anger | 0.930 | 0.6281/ 0.6281000 | | |
| Stress-related paranoid ideation | 21.368 | < 0.001 / < 0.001 | 0.266 | 0.6058/ 0.9151 |

Results of sequential analytic sub-steps n°2 and 3 used to determine the regression paths that link the reproduction-maintenance trade-off and BPD symptoms, and their variation between men and women.

[a] chi-square differences between a model with versus a model without the link of interest between the reproduction-maintenance trade-off and the individual BPD symptom of interest, using the DIFFTEST procedure (Asparouhov and Muthén 2010). A significant difference, highlighted in colour, indicates that there is a direct effect of the reproduction-maintenance trade-off on the probability of observing the BPD symptom.

[b] chi-square differences between (i) a model in which the direct effect of interest (of the reproduction-maintenance trade-off on the BPD symptom indicator) is constrained to be equal between men and women, and (ii) a model in which this direct effect is allowed to vary between men and women, while the other direct effects are constrained to be equal between men and women. A significant difference, highlighted in green, indicate that the direct effect of the reproduction-maintenance trade-off on BPD symptoms indicator, resulting from the previous analysis sub-step [a], varies between men and women.

**Table S10. Results of the alternative MIMIC model – Results of the sequential analytic steps used to determine the variation of the BPD symptoms thresholds between men and women.**

| | Test for the variation of the BPD symptoms thresholds between men and women | | Threshold [a] | |
|---|---|---|---|---|
| Symptom | χ2 difference [b] | p difference (Uncorr / Benjamini-Hochberg) | Male | Female |
| Frantic efforts to avoid real/imagined abandonment | 4.976 | 0.0257/ 0.04770000 | 1.837 | 1.930 |
| Unstable/intense interpersonal relationships | 3.466 | 0.0627/ 0.07053750 | 1.803 | 1.729 |
| Identity disturbance | 4.926 | 0.0265/ 0.04770000 | 2.054 | 1.956 |
| Impulsivity | 25.325 | < 0.001 / < 0.001 | 1.542 | 1.717 |
| Suicidal/self-mutilation behavior | 4.336 | 0.0373/ 0.04795714 | 2.516 | 2.373 |
| Affective instability | 4.928 | 0.0264/ 0.04770000 | 2.239 | 2.120 |

| | | | | |
|---|---|---|---|---|
| Chronic feelings of emptiness | 4.572 | 0.0325/ 0.04795714 | 2.183 | 2.064 |
| Inappropriate/intense anger | 9.452 | 0.0021/ 0.00945000 | 1.699 | 1.816 |
| Stress-related paranoid ideation | NA | NA | 2.027 | 1.981 |

[a] Thresholds correspond to the severity parameter (i.e. the threshold of the link function between BPD severity and BPD symptom) which corresponds to the threshold on latent BPD severity factor at which a symptom has a 50% probability of being endorsed.

[b] chi-square differences, using the DIFFTEST procedure (Asparouhov and Muthén 2010), between (i) a model in which the threshold for the BPD symptom of interest is constrained to be equal between men and women, and (ii) a model in which this threshold is allowed to vary between men and women, while the other thresholds as well as the slopes of the link function between BPD severity latent variable and BPD symptoms (i.e. the discrimination parameters) are constrained to be equal between men and women. A significant difference, highlighted in green, indicates that there is sex difference in the probability of observing the BPD symptom that does not depend on the reproduction-maintenance trade-off or the BPD severity. Consequently, a significant difference corresponds to a DIF between men and women that is not influenced by the reproduction-maintenance trade-off or the BPD severity, but by unknown factors not measured in our model.

### Table S11. Results of the alternative MIMIC model – Results of the sequential analytic steps used to determine the variation of the BPD severity indicators slopes between men and women.

| | Test for the variation of the BPD symptoms slopes between men and women | | Slope [a] | | |
|---|---|---|---|---|---|
| Symptom | χ2 difference [b] | p difference (Uncorr / Benjamini-Hochberg) | Fixed | Male | Female |
| Frantic efforts to avoid real/imagined abandonment | 0.303 | 0.5819/ 0.7716857 | 0.736 | 0.742 | 0.730 |
| Unstable/intense interpersonal relationships | 0.275 | 0.6002/ 0.7716857 | 0.741 | 0.735 | 0.745 |
| Identity disturbance | 0.051 | 0.8209/ 0.9235125 | 0.734 | 0.731 | 0.736 |
| Impulsivity | 1.595 | 0.8209/ 0.4790250 | 0.654 | 0.668 | 0.640 |
| Suicidal/self-mutilation behavior | 1.552 | 0.2129/ 0.4790250 | 0.616 | 0.584 | 0.631 |
| Affective instability | 6.684 | 0.0097/ 0.0873000 | 0.800 | 0.756 | 0.822 |

| | | | | | |
|---|---|---|---|---|---|
| Chronic feelings of emptiness | 4.039 | 0.0445/ 0.2002500 | 0.845 | 0.818 | 0.862 |
| Inappropriate/intense anger | 0.290 | 0.5904/ 0.7716857 | 0.773 | 0.767 | 0.777 |
| Stress-related paranoid ideation | NA | NA | 0.616 | 0.793 | 0.788 |

[a] Slope corresponds to the discrimination parameter (i.e., the slope of the link function between BPD severity and BPD symptom also known as the loading of the BPD symptom on BPD severity) which corresponds to the change in the probability of endorsing the symptom across increasing levels of the latent BPD severity factor. This means that there is variation in BPD severity that contribute to difference in the probability to observe this BPD symptom, above and beyond the effect of the reproduction-maintenance trade-off.

[b] chi-square differences, using the DIFFTEST procedure (Asparouhov and Muthén 2010), between (i) a model in which the slope for the BPD symptom of interest is constrained to be equal between men and women, and (ii) a model in which this slope is allowed to vary between men and women, while the threshold for other BPD symptoms are allowed to vary between men and women and the slope for other BPD symptoms are constrained to be equal between men and women. A significant difference would indicate that the effect of the BPD severity latent factor on the probability to observe a BPD symptom varies between men and women. A significant difference corresponds to another kind of DIF between men and women.

### Table S12. Results of the alternative MIMIC model – Parameter estimates of the final alternative MIMIC model

| **Alternative Final Multiple Indicator Multiple Cause (MIMIC) Male model** | | | | | | | |
|---|---|---|---|---|---|---|---|
| **Model part** | **Latent variables** | | **Indicators** | **Unstd. est.** | **se** | **p** | **Std. est.** |
| **Measurement model** | BPD severity | BY | Unstable/intense interpersonal relationships | 0.741 | 0.010 | < .001 | STDY 0.794 |
| | BPD severity | BY | Identity disturbance | 0.734 | 0.012 | < .001 | STDY 0.779 |
| | BPD severity | BY | Impulsivity | 0.654 | 0.013 | < .001 | STDY 0.701 |
| | BPD severity | BY | Suicidal/self-mutilation behaviour | 0.616 | 0.022 | < .001 | STDY 0.649 |
| | BPD severity | BY | Frantic efforts to avoid real/imagined abandonment | 0.736 | 0.010 | < .001 | STDY 0.786 |

| | | | | | | | |
|---|---|---|---|---|---|---|---|
| | BPD severity | BY | Inappropriate/intense anger | 0.773 | 0.010 | < .001 | STDY 0.823 |
| | BPD severity | BY | Stress-related paranoid ideation | 0.790 | 0.012 | < .001 | STDY 0.847 |
| | BPD severity | BY | Affective instability | 0.800 | 0.011 | < .001 | STDY 0.832 |
| | BPD severity | BY | Chronic feelings of emptiness | 0.845 | 0.012 | < .001 | STDY 0.890 |
| | Reproduction-maintenance trade-off | BY | Metabolic syndrome | 0.187 | 0.017 | < .001 | STDXY 0.202 |
| | Reproduction-maintenance trade-off | BY | Body Mass Index | 0.069 | 0.009 | < .001 | STDXY 0.146 |
| | Reproduction-maintenance trade-off | BY | Perceived physical health | -0.2712 | 0.016 | < .001 | STDXY -0.300 |
| | Reproduction-maintenance trade-off | BY | Age at first sexuel intercourse | -0.150 | 0.008 | < .001 | STDXY -0.433 |
| | Reproduction-maintenance trade-off | BY | History of sexually transmitted diseases | 0.293 | 0.071 | < .001 | STDY 0.308 |
| | Reproduction-maintenance trade-off | BY | Number of marriages | 0.148 | 0.012 | < .001 | STDXY 0.206 |
| | Reproduction-maintenance trade-off | BY | Number of children | 0.209 | 0.027 | < .001 | STDXY 0.124 |
| | Metabolic syndrome | WITH | Perceived physical health | -0.139 | 0.0009 | < .001 | STDXY -0.172 |
| | Metabolic syndrome | WITH | Body Mass Index | 0.260 | 0.005 | < .001 | STDXY 0.558 |
| | Body Mass Index | WITH | Perceived physical health | -0.042 | 0.004 | < .001 | STDXY -0.096 |
| | Number of marriages | WITH | Number of children | 0.337 | 0.008 | < .001 | STDXY 0.301 |
| **Structural model** | BPD severity | ON | Reproduction-maintenance trade-off | 0.339 | 0.025 | < .001 | STDXY 0.330 |

| | | | | | | | |
|---|---|---|---|---|---|---|---|
| | Impulsivity | ON | Reproduction-maintenance trade-off | 0.098 | 0.018 | < .001 | STDXY 0.102 |
| | Suicidal/self-mutilation behaviour | ON | Reproduction-maintenance trade-off | 0.138 | 0.025 | < .001 | STDXY 0.142 |
| | Chronic feelings of emptiness | ON | Reproduction-maintenance trade-off | -0.039 | 0.019 | 0.046 | STDXY -0.040 |
| | Stress-related paranoid ideation | ON | Reproduction-maintenance trade-off | -0.069 | 0.019 | < .001 | STDXY -0.072 |
| | BPD severity | ON | Early life adversity | 2.826 | 0.236 | < .001 | STDXY 0.193 |
| | Reproduction-maintenance trade-off | ON | Early life adversity | 5.177 | 0.288 | < .001 | STDXY 0.383 |
| **Thresholds** | | | Unstable/intense interpersonal relationships | 1.763 | 0.020 | < .001 | STDY 1.692 |
| | | | Identity disturbance | 2.053 | 0.032 | < .001 | STDY 1.956 |
| | | | Impulsivity | 1.541 | 0.025 | < .001 | STDY 1.478 |
| | | | Suicidal/self-mutilation behaviour | 2.518 | 0.058 | < .001 | STDY 2.389 |
| | | | Frantic efforts to avoid real/imagined abandonment | 1.837 | 0.032 | < .001 | STDY 1.757 |
| | | | Inappropriate/intense anger | 1.700 | 0.026 | < .001 | STDY 1.622 |
| | | | Stress-related paranoid ideation | 2.096 | 0.026 | < .001 | STDY 2.018 |
| | | | Affective instability | 2.239 | 0.041 | < .001 | STDY 2.126 |
| | | | Chronic feelings of emptiness | 2.183 | 0.042 | < .001 | STDY 2.075 |

| **Alternative Final Multiple Indicator Multiple Cause (MIMIC) Female model** | | | | | | | |
|---|---|---|---|---|---|---|---|
| **Model part** | **Latent variables** | | **Indicators** | **Unstd. est.** | **se** | **p** | **Std. est.** |

| | | | | | | | |
|---|---|---|---|---|---|---|---|
| **Measurement model** | BPD severity | BY | Unstable/intense interpersonal relationships | 0.741 | 0.010 | < .001 | STDY 0.787 |
| | BPD severity | BY | Identity disturbance | 0.734 | 0.012 | < .001 | STDY 0.786 |
| | BPD severity | BY | Impulsivity | 0.654 | 0.013 | < .001 | STDY 0.691 |
| | BPD severity | BY | Suicidal/self-mutilation behaviour | 0.616 | 0.022 | < .001 | STDY 0.635 |
| | BPD severity | BY | Frantic efforts to avoid real/imagined abandonment | 0.736 | 0.010 | < .001 | STDY 0.774 |
| | BPD severity | BY | Inappropriate/intense anger | 0.773 | 0.010 | < .001 | STDY 0.812 |
| | BPD severity | BY | Stress-related paranoid ideation | 0.790 | 0.012 | < .001 | STDY 0.841 |
| | BPD severity | BY | Affective instability | 0.800 | 0.011 | < .001 | STDY 0.830 |
| | BPD severity | BY | Chronic feelings of emptiness | 0.845 | 0.012 | < .001 | STDY 0.891 |
| | Reproduction-maintenance trade-off | BY | Metabolic syndrome | 0.146 | 0.010 | < .001 | STDXY 0.167 |
| | Reproduction-maintenance trade-off | BY | Body Mass Index | 0.093 | 0.007 | < .001 | STDXY 0.167 |
| | Reproduction-maintenance trade-off | BY | Perceived physical health | -0.232 | 0.011 | < .001 | STDXY -0.244 |
| | Reproduction-maintenance trade-off | BY | Age at first sexuel intercourse | -0.177 | 0.005 | < .001 | STDXY -0.532 |
| | Reproduction-maintenance trade-off | BY | History of sexually transmitted diseases | 0.347 | 0.063 | < .001 | STDY 0.360 |
| | Reproduction-maintenance trade-off | BY | Number of marriages | 0.175 | 0.008 | < .001 | STDXY 0.263 |
| | Reproduction-maintenance trade-off | BY | Number of children | 0.427 | 0.019 | < .001 | STDXY 0.258 |
| | Metabolic syndrome | WITH | Perceived physical health | -0.213 | 0.008 | < .001 | STDXY -0.244 |

| | | | | | | | |
|---|---|---|---|---|---|---|---|
| | Metabolic syndrome | WITH | Body Mass Index | 0.334 | 0.005 | < .001 | STDXY 0.588 |
| | Body Mass Index | WITH | Perceived physical health | -0.115 | 0.005 | < .001 | STDXY -0.192 |
| | Number of marriages | WITH | Number of children | 0.149 | 0.008 | < .001 | STDXY 0.126 |
| **Structural model** | BPD severity | ON | Reproduction-maintenance trade-off | 0.319 | 0.024 | < .001 | STDXY 0.347 |
| | Impulsivity | ON | Reproduction-maintenance trade-off | 0.097 | 0.016 | < .001 | STDXY 0.106 |
| | Suicidal/self-mutilation behaviour | ON | Reproduction-maintenance trade-off | 0.146 | 0.025 | < .001 | STDXY 0.145 |
| | Chronic feelings of emptiness | ON | Reproduction-maintenance trade-off | -0.027 | 0.020 | 0.179 | STDXY -0.041 |
| | Stress-related paranoid ideation | ON | Reproduction-maintenance trade-off | -0.066 | 0.020 | 0.001 | STDXY -0.075 |
| | BPD severity | ON | Early life adversity | 1.958 | 0.177 | < .001 | STDXY 0.168 |
| | Reproduction-maintenance trade-off | ON | Early life adversity | 5.462 | 0.180 | < .001 | STDXY 0.489 |
| **Thresholds** | | | Unstable/intense interpersonal relationships | 1.763 | 0.020 | < .001 | STDY 1.670 |
| | | | Identity disturbance | 1.959 | 0.030 | < .001 | STDY 1.868 |
| | | | Impulsivity | 1.719 | 0.024 | < .001 | STDY 1.616 |
| | | | Suicidal/self-mutilation behaviour | 2.371 | 0.039 | < .001 | STDY 2.179 |
| | | | Frantic efforts to avoid real/imagined abandonment | 1.931 | 0.027 | < .001 | STDY 1.809 |
| | | | Inappropriate/intense anger | 1.817 | 0.028 | < .001 | STDY 1.701 |
| | | | Stress-related paranoid ideation | 2.097 | 0.026 | < .001 | STDY 1.989 |

| | | | | |
|---|---|---|---|---|
| Affective instability | 2.103 | 0.038 | < .001 | STDY 1.959 |
| Chronic feelings of emptiness | 2.064 | 0.037 | < .001 | STDY 1.939 |

Elements of the Mplus model syntax (L. K. Muthén & Muthén, 2017): BY can be read as "is measured by"; ON can be read as “is regressed on”; WITH can be read as “is correlated with”.
STDY corresponds to estimates standardization with respect to the latent dependent variable which is suitable for binary variables. The STDY standardized estimate is interpreted as the change in y in y standard deviation units when x changes from zero to one. The STDXY standardized estimate is interpreted as the change in y in y standard deviation units for a standard deviation change in x. Thresholds correspond to the severity parameter (i.e., the threshold of the link function between BPD severity and BPD symptom) which corresponds to the threshold on latent BPD severity factor at which a symptom has a 50% probability of being endorsed.

**Figure S2. Final alternative Multiple Indicator Multiple Cause (MIMIC) multi-group model in Men vs Women.**

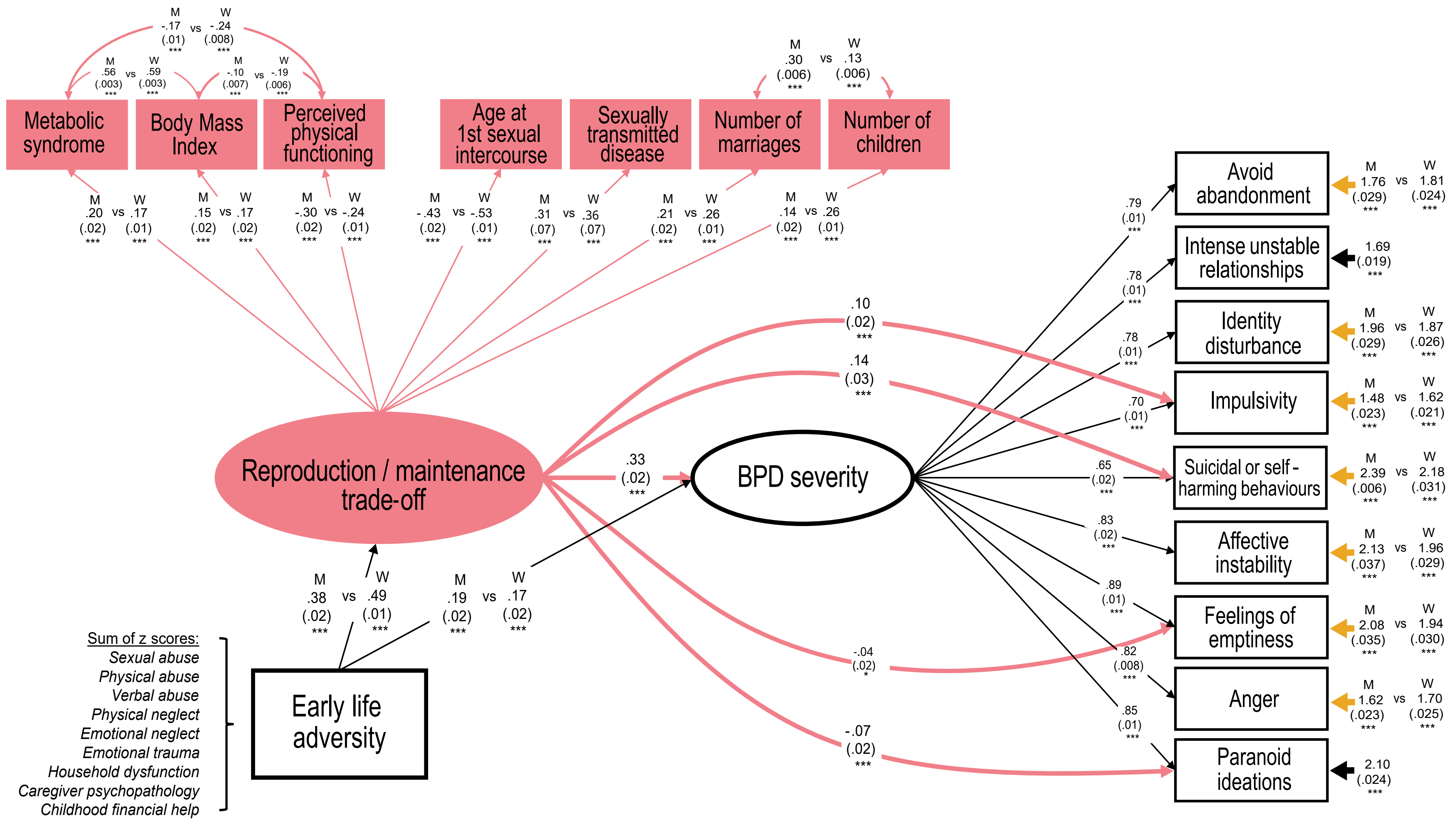

Simplified representation of final MIMIC model which showed a good fit (CFI= 0.982, TLI= 0.975, RMSEA= 0.016[ 0.015–0.017], SRMR=0.041) Ellipses represent latent constructs, rectangles represent observed indicators. Paths between indicators and the reproduction/maintenance trade-off latent factor represent factor loadings. Paths between BPD symptoms indicators and the BPD severity latent factor also represent factor loadings. Paths between the reproduction/maintenance trade-off latent factor and BPD symptoms represent direct DIF. Dashed lines represent paths that are not statistically significant. The arrows to the right of the BPD severity indicators represent the thresholds of the link function between BPD severity and BPD symptom. The yellow arrows indicate that there is a significant sex difference in the probability of observing the BPD symptom that does not depend on the reproduction-maintenance trade-off or the BPD severity. Consequently, a yellow arrow indicates that there is a DIF between men and women that is not influenced by the reproduction-maintenance trade-off or the BPD severity, but by unknown factors not measured in our model. The letter ‘M’ indicates results of the full MIMIC model in the Men subsample. The letter ‘W’ indicates results of the full MIMIC model in the Women subsample. Significance codes: n.s. for p>0.05, * for p<0.05, ** for p<0.01, *** for p<0.001.

### Method S6. 10-FOLD CROSS VALIDATION

To test the capacity of our main model to generalize its predictions to out-of-sample data, we employed a 10-fold cross-validation procedure, following three main steps:

1. The full data set is randomly partitioned into 10 folds of nearly equal size.

2. Subsequently, 10 iterations of training and validation are performed such that within each iteration a different fold of the data is held-out for validation (i.e. the test data, here representing 10% of the whole sample) while the remaining k-1 folds are used for fitting (i.e. the training data, here representing 90% of the whole sample).

3. At each iteration, the model is fit on the training data and has its parameters fixed to these results.

Cross-validation performance was indexed by the global fit indices (RMSEA, CFI, TLI, SRMR statistics) which quantifies the adequacy of the fitted model to the test data (see Table S13 right below). The overall stability of these measures across the multiple partitioning of the dataset were used as indicators of the robustness of the results to sampling variability.

### Table S13. 10-fold cross validation

| Fold | X2 (train/test) | RMSEA (train/test) | CFI (train/test) | TLI (train/test) | SRMR (train/test) |
|---|---|---|---|---|---|
| 1 | 1070.772/451.259 | 0.017 (0.016-0.018)/0.026 (0.023-0.030) | 0.985/0.961 | 0.979/ 0.945 | 0.038/0.077 |
| 2 | 1071.136/310.799 | 0.017(0.016 -0.018)/0.017(0.012- 0.021) | 0.985/0.986 | 0.979/0.980 | 0.039/0.059 |
| 3 | 1083.692/ 302.319 | 0.017(0.016-0.018)/0.020(0.016-0.024) | 0.985/0.981 | 0.980/0.973 | 0.040/0.047 |
| 4 | 1067.544/332.392 | 0.017 (0.016-0.018)/0.019(0.015-0.023) | 0.985/0.984 | 0.979/0.977 | 0.040/0.062 |
| 5 | 1056.614/403.280 | 0.017(0.016-0.018)/0.024(0.020-0.027) | 0.985/0.972 | 0.979/0.960 | 0.039/0.079 |
| 6 | 1035.556/389.667 | 0.016(0.015-0.018)/0.023(0.019-0.026) | 0.986/0.971 | 0.980/0.960 | 0.039/0.067 |
| 7 | 1052.509/401.954 | 0.017(0.016-0.018)/0.024(0.020-0.027) | 0.986/0.968 | 0.980/0.955 | 0.037/0.081 |
| 8 | 1112.372/292.255 | 0.017(0.016-0.018)/0.015(0.010-0.019) | 0.984/0.989 | 0.978/0.984 | 0.041/0.060 |
| 9 | 1044.618/366.184 | 0.017(0.016 -0.018)/0.021(0.017 -0.025) | 0.985/0.981 | 0.979/0.973 | 0.038/0.052 |
| 10 | 1027.119/498.619 | 0.016(0.015 - 0.017)/0.029(0.026-0.032) | 0.986/0.957 | 0.980/0.940 | 0.039/0.074 |
